# Digitizing the coral reef: machine learning of underwater spectral images enables dense taxonomic mapping of benthic habitats

**DOI:** 10.1101/2022.03.28.485758

**Authors:** Daniel Schürholz, Arjun Chennu

## Abstract

1. Coral reefs are the most biodiverse marine ecosystems, and host a wide range of taxonomic diversity in a complex spatial habitat structure. Existing coral reef survey methods struggle to accurately capture the taxonomic detail within the complex spatial structure of benthic communities.
2. We propose a workflow to leverage underwater hyperspectral transects and two machine learning algorithms to produce dense habitat maps of 1150 m^2^ of reefs across the Curaçao coastline. Our multi-method workflow labelled all 500+ million pixels with one of 43 classes at taxonomic family, genus or species level for corals, algae, sponges, or to substrate labels such as sediment, turf algae and cyanobacterial mats.
3. With low annotation effort (2% pixels) and no external data, our workflow enables accurate (Fbeta 87%) survey-scale mapping, with unprecedented thematic and spatial detail. Our assessments of the composition and configuration of the benthic communities of 23 transect showed high consistency.
4. Digitizing the reef habitat structure enables validation and novel analysis of pattern and scale in coral reef ecology. Our dense habitat maps reveal the inadequacies of point sampling methods to accurately describe reef benthic communities.

## Introduction

Under rapidly changing environmental conditions, the need for accurate and speedy ecological assessment of marine and freshwater ecosystems has greatly increased. This is particularly pressing for coral reefs, which are the most biologically diverse marine ecosystems on the planet, but have suffered significant deterioration in recent years due to a variety of stressors (Hughes et al., 2020). The continuing deterioration of coral reef health worldwide will endanger many of the ecosystems services that these reefs provide to coastal populations and other associated systems (Hoegh-Guldberg et al., 2017). Furthermore, this long-term degradation of reefs confounds an inter-generational understanding of baseline reef health that informs reef restoration and management interventions (Muldrow et al., 2020), thus highlighting the need for objective assessments of reef health through monitoring.

Modern reef monitoring efforts focus on the creation of benthic habitat maps, as they capture the spatial distributions of species, habitat features and their environmental context (Guisan et al., 2013; Roelfsema et al., 2020). Such information captured over a long time series forms the basis of scientific evaluation of the ecosystem’s evolution, and underpins the decisions for management, conservation and restoration (Foo & Asner, 2019). The spatial, temporal and thematic scales of ecosystem mapping have a critical influence on the viability of specific scientific analyses (Lecours, 2017), such as elucidating functional drivers, detecting community phase shifts or signalling deterioration of habitats. Most reef inventories compiled from in-situ surveys lack sufficient taxonomic and spatial detail, and have been carried out in only 0.01% – 0.1% of coral reef regions worldwide (Hochberg & Gierach, 2021). Also, many surveys do not report any uncertainty information which limits the utility of the data for scaling up to ecosystem-level studies (Reverter et al., 2022). Thus, a priority for future in-situ reef surveys should be wider biogeographic coverage, clearer estimates of uncertainty, and greater taxonomic and spatial detail at the survey scale.

Satellites are increasingly used to map shallow benthic habitats and analyse regional and global phenomena affecting coral reefs (Hedley et al., 2016; Heron et al., 2016). With recent enhanced spectral and spatial resolutions of remotely sensed images (0.5–10 m per pixel), better reef monitoring products have been enabled, such as geomorphological zonation of reefs (Kennedy et al., 2021) and benthic habitat maps (B. Lyons et al., 2020; Roelfsema et al., 2021). However, validating the accuracy of satellite-derived maps is a difficult task, impeded by the lack of in-situ validation datasets and the lack of error estimation in those that do exist (Hochberg & Gierach, 2021; Phinn et al., 2012). While remote sensing offers a viable approach for large scale analyses of reefs, current satellite sensors lack spatial resolution to represent small organisms (<0.5m) and the spectral resolution to potentially differentiate organisms to a deep taxonomic description (Hochberg et al., 2003; Muller-Karger et al., 2018). In contrast, in-situ surveys can provide enhanced spatial and spectral resolutions in underwater imagery, made available by advancements in instrumentation (Chennu et al., 2017), and robotic platforms for both aerial (Casella et al., 2017) and underwater (Armstrong et al., 2019) usage. Improvements in machine learning (ML), especially the application of artificial neural networks, have contributed to better accuracy and throughput of efforts in automated classification (Beijbom et al., 2015; González-Rivero et al., 2020) and semantic segmentation (Alonso et al., 2019) of benthic images. Images acquired through underwater/proximal sensing, and mapped through scalable and automated workflows, can provide a consistent source of validation for current and upcoming ecosystem-level studies.

Deriving validation support from in-situ surveys requires careful design conformity between the proximal and remote sensing campaigns (Roelfsema & Phinn, 2013). For example, the lack of conformity in the set of labels used between satellite and in-situ studies is a major confounding factor (Foody, 2004). The labelspace of global maps usually include broad functional groups in the reef (coral, algae, sediment, etc.), some status indicators (dead, alive, bleached) or morphological descriptions (branching, massive, weedy) (Kennedy et al., 2021). This multi-faceted and easy-to-interpret view of the reef structure is useful for coastal management (Roelfsema et al., 2020). In contrast, in-situ studies achieve thematic scales that identify organisms down to genus or species level, as well as different substrata (sand, rock, rubble) and the substrate-associated communities (cyanobacterial mats, turf algae) to render a detailed view of the biotic and abiotic components (Althaus et al., 2015). Capturing habitat structure with a detailed labelspace is typically limited by the availability of expert time or by logistical constraints. For this reason, reef habitat structure is severely undersampled – both spatially and thematically – in a majority of reef studies.

Here we demonstrate how to produce accurate, dense and detailed maps of coral reef habitats from underwater surveys (Fig 1). Dense means that all regions/pixels in each transect are assigned a (biotic or abiotic) habitat label, resulting in full semantic segmentation of the transect without any catch-all ‘background’ labels. Detailed refers to the thematic detail that is captured by the labels, such as either being taxonomic (species, genus, etc) or functional (corals, sponges, etc). We leverage machine learning of underwater hyperspectral transects captured over multiple weeks and locations along the coast of the Caribbean island of Curaçao (Chennu et al., 2017; Rashid & Chennu, 2020). Our workflow description (Fig 1) considers all the steps from the field survey to the classification of 500+ million pixels to the validation of aggregate habitat descriptions derived from the detailed habitat maps. We show how dense maps can be produced, with clear assessments of accuracy, into multiple thematic scales, either at a functional (“reefgroups”) or taxonomic (“detailed”) labelspace. By implementing two independent ML methods in parallel, we show that the machine-produced habitat maps are consistent in terms of habitat descriptors, and can be generated with workflows of different complexities. These ML methods can be used to map and analyse spectral transects at the survey scale, without the need to source training data from external datasets. We provide an assessment of the consistency between the habitat structure of the benthic community as described by the maps produced with each ML method. Finally, we exploit the dense habitat maps of island-wide transects to reveal the inadequacies of point sampling methods to accurately describe reef benthic communities.

**Fig. 1.**
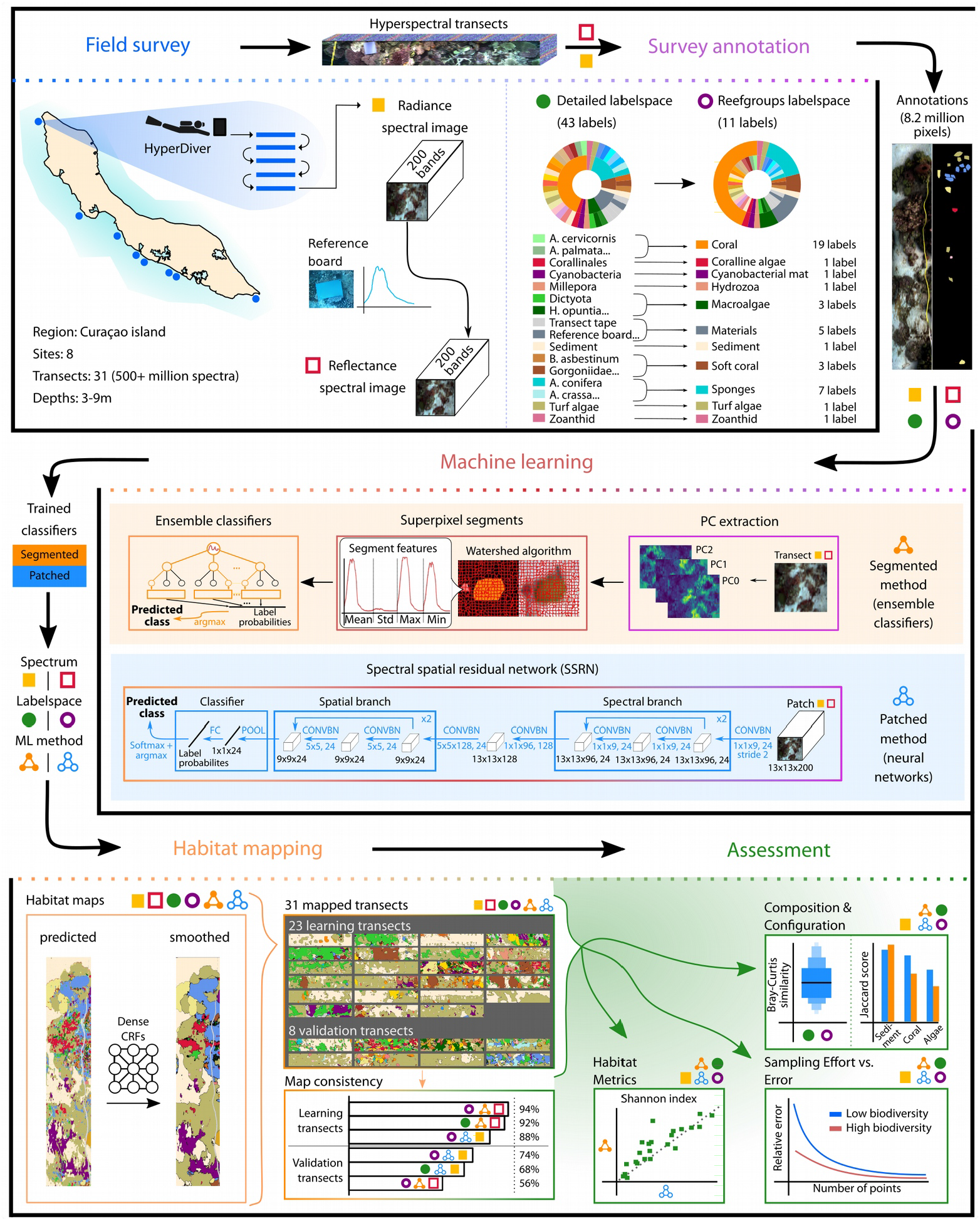
Schematic of scalable survey-mapping-assessment workflow for digitizing reef habitats. The underwater survey with the HyperDiver at 8 study sites over 147 transects (50 m each) in Curacao produced radiance and reflectance spectral images. A subset of 31 transect images were annotated into the detailed (43 classes) labelspace and aggregated into the reefgroups (11 classes) labelspace. Annotated regions from 23 transects were used in two separate machine learning methods to classify each region of the spectral images into each labelspace. The segmented method used ensemble learning on image superpixel features, while the patched method used spatial-spectral neural network learning of image patches. The classifiers were used to predict the label probabilities at each of 500+ million pixels in all 31 transects. The classifier-predicted probability maps were smoothed and converted into densely-labeled smooth habitat maps. The habitat maps were assessed for their consistency with reference annotations as well as their ability to describe the composition and configuration of the benthic communities in the transects. The effort-vs-error relationship for point-count sampling of the reef habitats was assessed using virtual sampling of the dense habitat maps.

## Materials and Methods

### Underwater hyperspectral surveying

Underwater hyperspectral transects were acquired with the HyperDiver surveying system (Chennu et al., 2017), which contains a line imager that captures hyperspectral images of the seafloor. Hyperspectral transects were acquired in a survey conducted along the leeward coastline of the Caribbean island of Curaçao (Fig 1 “Field Survey”). At each of the eight survey sites, 10 to 20 transect images of 50 m long were acquired by divers at varying depths (3 m to 9 m range). The resulting dataset contains 147 hyperspectral transects, from which 31 transects were selected for testing the proposed workflow. Each of the 640 pixels in the line-of-view contained 12-bits of radiometric information for each of the 480 wavelengths bands in the 400-800 nm range. The spectral images were interpolated and reduced to 200 bands of 8-bit radiometric precision. Each of the transect images was captured under natural and varying light conditions due to depth, cloud cover, water surface conditions, etc. Each transect’s radiance image was converted to pseudo-reflectance images by dividing out the average radiance signal of a gray reference board present in each transect scene. A detailed description of the acquisition and processing is available in a data descriptor (Rashid & Chennu, 2020).

### Benthic annotation and thematic flexibility

Annotations were created by human experts to support automated classification of the transects by machine learning methods. The annotations consisted of 2089 small polygons covering 8.2 million pixels with a corresponding habitat label across the 31 transects (Fig S1, S2; Fig 1 “Survey annotation”). Biotic classes were annotated to the deepest taxonomic level possible, such as family, genus or species. Substrate classes were represented as sediment, cyanobacterial mat or turf algae. Survey materials were also included to give semantic labels to any object found in the transects, i.e. transect tape or reference board. Three classes were removed given their very low representation in the selected transects (<2 annotated regions). The resulting “detailed” labelspace had 43 final labels (Fig S3). Loosely speaking, the detailed labelspace describes the habitat in the perspective of a reef ecologist.

To describe the habitat and community from the perspective of a reef manager, each label in the detailed labelspace was then abstracted to a corresponding functional reef group class (Fig SS1). As an example, the 19 coral species and genera were abstracted to a class called “Coral”. The 11 resulting classes formed the “reefgroups” labelspace. This thematic flexibility allowed us to run the ML setup with annotations in either labelspace, to measure the workflow’s ability to classify into both labelspaces correctly. To compare classifications across labelspaces, we created an abstracted “detailed-to-reefgroups” version of the detailed lablespace maps, that is, assigning all labels to their corresponding reefgroups label. Then the reef community composition was compared between the detailed-to-reefgroups maps and the reefgroups maps (Fig 6E-F).

The reference annotations were created with a bias towards reducing human effort rather than providing uniform coverage of samples across the survey data (Rashid & Chennu, 2020). This resulted in a ML dataset with a relatively high degree of label imbalance, both when considered as a set of polygons or as a set of pixels across the annotated transects (Fig S1, S2). The degree of imbalance meant that, for example, the 5 most abundant classes (*Sediment, Turf algae, D. strigosa, Dictyota, S. siderea*) had 789 polygons and 3.16 million pixels, while the 5 rarest classes (*A. cauliformis, B. asbestinum, D. stokesii, Zoanthid, L. variegata*) had only 14 polygons and 18464 pixels.

The annotated data consisted of 23 “learning transects” (373+ million pixels), that were used to train and test classifiers in the machine learning steps of our workflow, and 8 separate “validation transects” (150+ million pixels) that were used to assess the performance of our workflow on data fully outside the learning transects. Overall, it was possible to represent each annotated transect with two types of signal (radiance or reflectance) and labelspaces (detailed or reefgroups) for machine learning towards automated classification.

### Machine learning for benthic mapping

Machine learning classifiers were created to predict the identity of each image region based on its spectral-spatial features (Fig 1 “Machine learning”). Two separate ML methods — “patched” and “segmented” — were independently implemented for each combination of signal type (radiance, reflectance) and labelspace (detailed, reefgroups)l. The prediction of each method for each image region was a probability value for each label/class in a labelspace.

For the patched method, a deep learning network called spectral-spatial residual network was used to train a classifier (Zhong et al., 2018). This network identifies spectral and spatial features by first convolving 1D filters in a spectral branch and then convolving 2D filters in a spatial branch over square patches from the hyperspectral image. Our hyperparameter tuning experiments indicated good performance for parameter values close to original study (see *Supplement*). For each pixel in the image, the probability of labels is predicted for the central pixel based on the neighbouring pixels in a regular image patch. The image was padded with reflection of border pixels to enable selection of patches at the image edges. After training, each transect was mapped by passing every image patch through the trained network to obtain the predicted label probabilities for the central pixel.

The segmented method consisted of three sequential processes to obtain the label probabilities for each image region (Fig 1). The first step was to reduce each transect image to six principal components and calculate the mean at each pixel. The second step was to segment the mean image into superpixels, which were contiguous irregularly-shaped image segments of similar pixels. The parameters for the watershed algorithm were a batch size of 2000 x 640 pixels, 12000 markers per batch and a compactness of 1000. This reduced the transect image into a set of irregularly shaped superpixels which over-segmented the objects in the image. For each hyperspectral image segment, descriptors were calculated for each spectral band: mean, standard deviation, minimum and maximum values, and concatenated into a vector of 800 features. These vectors were then used as input samples for a random forest ensemble classifier. The parameters for the classifier used for transect mapping were 300 base estimators, 2 minimum leaves per tree, 25 as the maximum tree depth and a minimum of 3 samples per split. The function used to measure the quality of a split was the Gini inequality function and the class weights were adjusted inversely to the class abundance in the samples. The classifier output was the label probabilities for an image segment, which were assigned to all the pixels within the segment to generate the class probability map of each transect.

We studied how the performanc of both classifiers depended on the quantity and quality of annotated spectral pixels (Fig 3A,B). The quantity of annotated data was measured as number of unique pixels used during training. Classifier performance was measured by training on various quantities of data but keeping the amount of computing effort constant (see *Supplement*). Furthermore, to measure the effect of the quality of the spectral information on the classifiers’ performance, a subset of equally-spaced spectral bands were selected (N=[10, 25, 50, 100, 150] out of 200) from the transect images (Fig 3C). Each method was separately trained and tested on the transect images with the subsampled spectral bands.

The class probability maps of each transects from either ML method was smoothed by refining the probabilities with dense conditional random fields (DCRF) (Krähenbühl & Koltun, 2011). DCRFs were used to update the probability value of each pixel based on the surrounding label probabilities. DCRFs interconnect every pixel in an image through a graph model, thus allowing fusing of long-range and short-range context within the image. The class probability maps were used as unary potential inputs to the DCRF, to obtain the smoothed probability maps (Fig 1 “Habitat mapping”; Fig 4A,B).

Each class probability map — from either ML method and with or without smoothing — was converted to a class map by assigning to each pixel the identity of the class with the highest probability. The result was a categorical habitat map where every pixel was assigned to one label in the labelspace of the trained classifier.

### Comparison of habitat structure

We assessed and compared the compositional and configurational structures of the reef benthos from the 23 (learning) transects distributed across Curaçao island (Fig 1 “Assessment”; Fig 6). Since both ML methods independently generated habitat maps in each labelspace of each transect, the habitat maps derived from each method were used for pairwise comparisons of the transect’s habitat structures. For each transect the percentage cover ( *P_i_*) was calculated as:

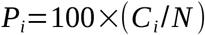

where *C_i_* is the count of pixels of class *i* in the transect and *N* is the total pixel count in the transect.

As a diversity measure for each transect we used Shannon diversity index (*H*′), defined as:

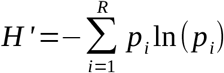

where *p_i_* is the proportion of elements of a class *i* and *R* is the total number of classes in the labelspace. To identify biases in the ML methods, we applied a Bland-Altman analysis on the habitat metrics derived for the transects from each ML method (Fig 6A-D). This analysis consists of two plots that help identify agreement between two quantitative methods of measurement. The first plots the values of both methods for the specific variable – percentage cover of a class or Shannon diversity index – against each other, to identify values deviating from the one-to-one correlation line. In the second plot the differences of the paired measurements are plotted against their averages, to identify the mean of the difference and its ±1.96 standard deviation lines. The bias is read as the gap between the mean of the difference and the 0 difference line. The two methods are considered to be in agreement if 95% of the values lie within the standard deviation lines in the second plot.

To measure the compositional similarity between two habitat maps we used the Bray-Curtis similarity (BCS) index defined as:

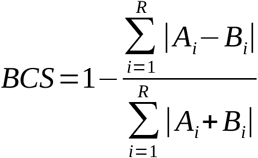

where *A_i_* and *B_i_* are counts of class *i* in the two maps. *R* is the total number of classes in the labelspace. The closer the BCS value is to 1 the more similar the composition of the two compared communities.

We measured the similarity in the configuration of the communities in two habitat maps by calculating the Jaccard score for each reefgroups class. The Jaccard score (*J*) is defined as:

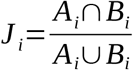

Where *A_i_* is the set of pixels of class *i* in map *A* and *B_i_* is the sets of pixels of class *i* in map *B*. When the Jaccard score is 1, then the two maps are identical in configuration and when the score is 0 then the maps are entirely dissimilar. A high Jaccard score across the habitat maps from both ML methods requires that both the identity and the location of the pixels are a match. Thus, it is a stringent measure of the similarity between two maps.

### Effort-vs-error of point-count sampling

We conducted simulations to estimate the error associated with a certain level of sampling effort in assessing the diversity and coverage of key groups through sparse point sampling of transects. For this experiment, we selected 4 transects with Shannon diversity index from low to high (H’ = {0.61, 1.26, 1.6, 2.64}). Each transect was divided into 50 non-overlapping quadrats of size 640 pixels × 406-705 pixels. Each of these quadrats was sparsely sampled with N = {5, 10, 20, 40, 80, 160, 240, 320, 480, 640, 960} randomly selected points. For example, this means that when N = 5 points, 250 points (50 quadrats x 5 points) were selected from the habitat map of the transect. The random sampling of the quadrats was conducted 250 times for each effort level. From each set of subsampled points in each transect, the coral coverage, sediment coverage and Shannon diversity index were calculated. For each metric, the relative deviation of the value obtained from the subsampled points from the value obtained from all the points in each transect was calculated. We selected 5% relative error as the limit for acceptable error The resulting error from changing sampling effort was compared for each of the transects containing a significantly different species diversity and coverage distribution (Fig 7).

## Results

### Automated workflow for scalable benthic mapping

Our workflow successfully employed two different ML methods to classify the transect images into habitat maps (Fig 1). The “segmented” method which used ensemble classifiers on irregular image segments generated through unsupervised segmentation, whereas the “patched” method used convolutional neural networks on regular image patches to produce predictions. Both methods were set up to take as input an image segment/patch from either of the signal types and produce the same output: an array of probability values for each label in the labelspace. The predictive performance of the trained classifiers was tested on a set of image annotations, which was spatially disjoint (no shared pixels) from the annotations used for training, as recommended in recent reviews (Paoletti et al., 2019). This testing set comprised of 15496 image segments for the segmented method and 50000 pixels for the patched method. The ML setup allowed us to combine the low-complexity segmented method (ensemble classifiers) and the high-complexity patched method (deep learning) as interchangeable ML components in the workflow for scalable reef mapping.

For the detailed labelspace, the segmented+reflectance combination had the best predictive performance with an Fbeta score of 84.5% (Fig 2A; Table S1). The classifier had 80% to 96% recall for a majority of the 43 labels with sufficient data support (see diagonal of Fig 2A). Some labels with low data support showed excellent recall (*A. archeri, A. crassa, B. asbestinum* and *D. stokesii*), while others showed significant errors (*Zoanthid, L. variegata* and *D. anchorata*). Despite having to distinguish between 43 labels, the segmented+reflectance classifier had a high Cohen’s kappa coefficient indicating performance which is 83.5% of a perfect classifier (Fig 2A; Table S1). For the same labelspace, the patched+radiance combination (Fig 2B; Fig S4) performed with a 9% lesser Fbeta score (76.7%). Most classes had a recall level between 70% and 90%, with some rare classes, such as *A. cauliformis* and *B. asbestinum,* having significant errors.

**Fig. 2.**
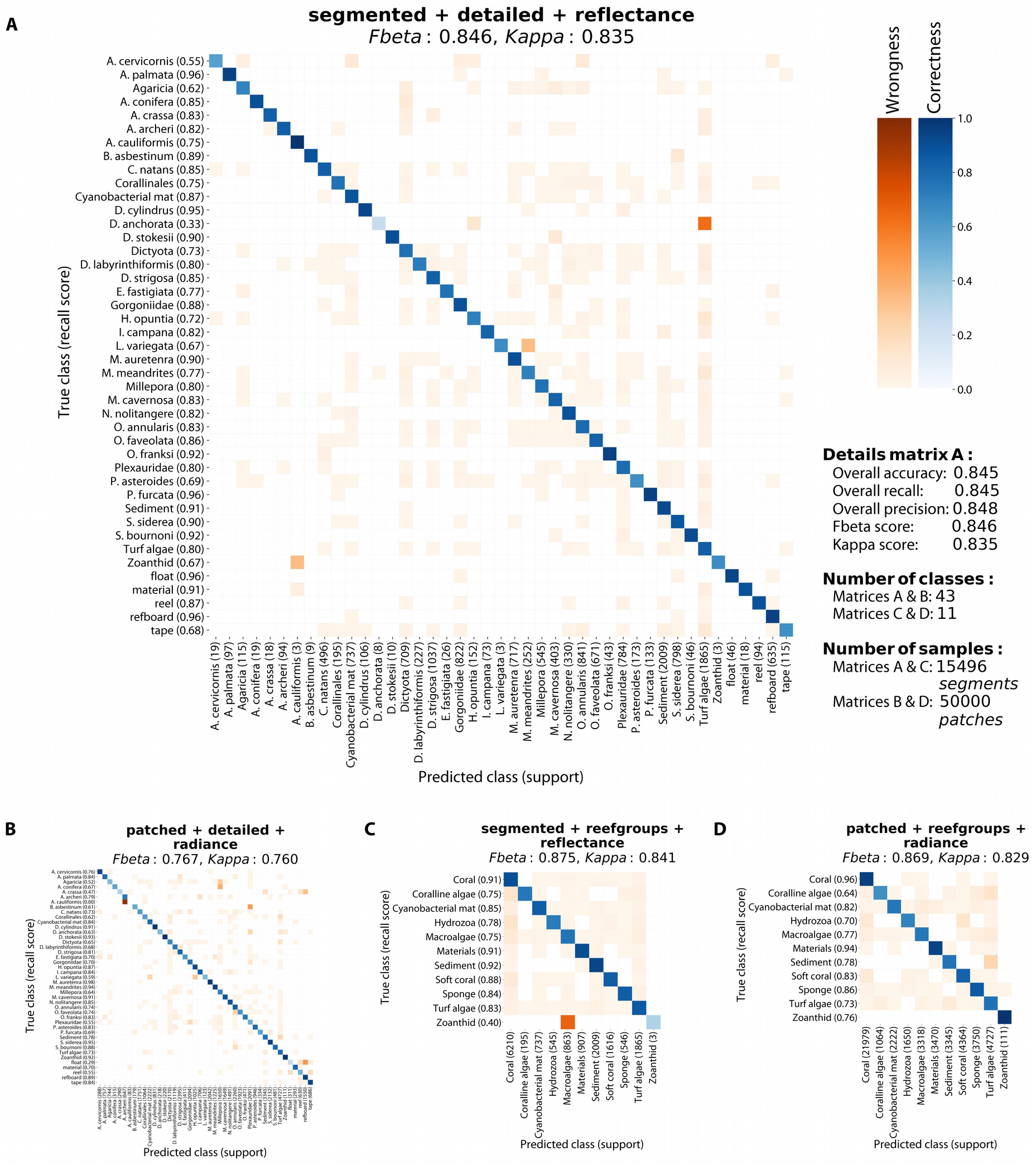
Performance evaluation of classifiers in detailed and reefgroup labelspaces. Confusion matrices of classifiers from a subset of ML method+labelspace+signal type combinations were used to asses performance on a held-out testing set. **(A)** The Fbeta score and other metrics (see side notes) showed excellent performance at 84.6%. There was generally little (<3%) to minor (<20%) confusions for the rare classes such as *Zoanthid* and *D. anchorata.* **(B)** There was little (<3%) to minor confusions (<20%) with rare classes such as *A. cauliformis* and *refboard.* **(C)** Some relevant confusions were found between the *Zoanthid* and *Macroalgae* classes. **(D)** Corals were classified with a recall of 96%.

For the reefgroups labelspace, the best predictive performance was 87.5% in the segmented+reflectance combination (Fig 2C) followed closely by the patched+radiance combination with 86.9% (Fig 2D). The latter showed high recall values for all 11 classes, even reaching 96% correctness for the *Coral* class – which consists of 19 different coral genera and species (Fig S3). The highest confusion occurred between *Turf algae* and *Sediment* classes. For both ML methods classifiers had a Cohen’s kappa score of 83% towards a perfect classifier (Fig 2; Table S1). Overall, both ML methods showed Fbeta scores between 72% to 87% with better performances on the reefgroups labelspace than on the detailed labelspace (Fig 2; Fig S4-S8; Table S1).

Both ML methods had different responses to the amount of unique input data seen during training. The classifier performance in the segmented method improved significantly with greater quantities of training data (Fig 3A). The greatest improvement was an Fbeta from 76% with 1.47 million pixels to 86% with 5.89 million pixels seen in the detailed labelspace using reflectance spectra. The performance also improved with greater quantities of radiance spectra, but overall the segmented method performed better on reflectance rather than radiance spectra. In contrast, the patched method showed no improvement with larger quantities of training data (Fig 3B). Instead, the performance in the detailed labelspace showed a 1% deterioration when the same computing effort was distributed over all the available data (5.93 million pixels). This seemingly unexpected result of poorer performance with more data can be understood by considering the lower number of training iterations over the larger dataset in order to maintain the same computing effort.

**Fig. 3.**
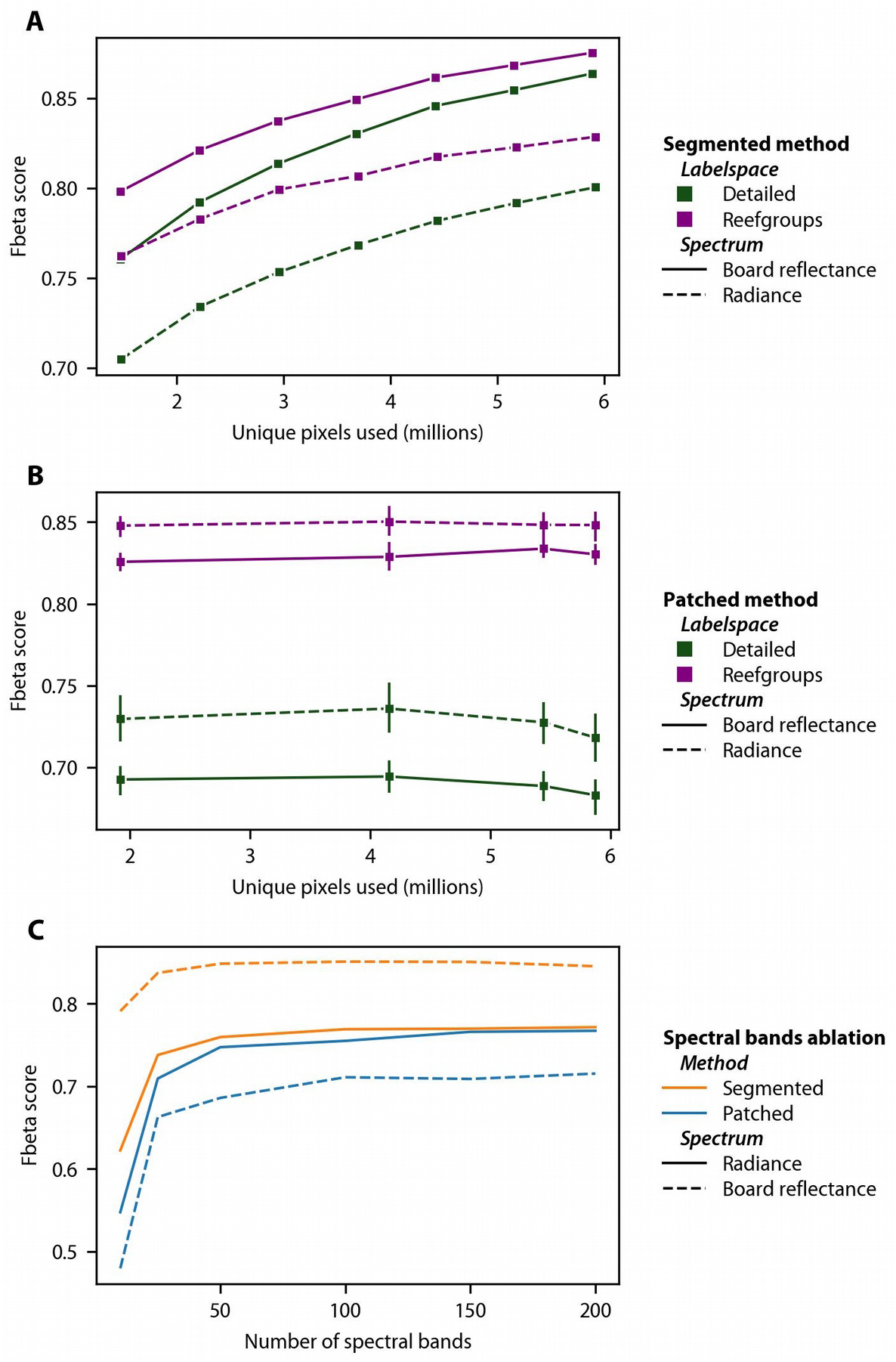
Effect of quantity and quality of spectral data on classifier performance. The effects of increasing the unique number of pixels in the data, but training with same computational effort, was measured (Fbeta score) for both methods and labelspaces. The segmented method **(A)** showed improved performance whereas the patched method **(B)** showed little change in performance. Both methods performed better at predicting into the reefgroups labelspace (with 11 classes) than the detailed labelspace (with 43 classes) irrespective of the spectral signal type. **(C)** Both methods performed better when using increasing number of (uniformly sampled) spectral bands for training, with limited improvement beyond 50 spectral bands.

The impact of data quality, or spectral resolution, on predictive performance was assessed by using a subset of 10-100 spectral bands in the training data. In both the segmented and patched methods (Fig 3C), the predictive performance showed a strong 5% to 15% improvement when the number of spectral bands was increased from 10 to 25 with diminishing improvements when using 50+ spectral bands. Overall, the availability of greater spectral resolution, even when the bands were chosen without special consideration, had a large effect on the performance of the ML methods. To produce habitat maps for further analysis, classifiers that were trained on all 200 bands and with 62500 patches (~4.1 million unique pixels) and 62332 segments (~5.8 million unique pixels).

### Smooth habitat maps to digitize reef habitat structure

The label probability map obtained directly from the classifier’s prediction showed generally noisy spatial distribution, with many areas of low confidence (Fig 4). This effect of low and noisy confidence was larger in the detailed than in the reefgroups labelspace. Processing the predicted probability map with DCRFs produced a more uniform map of probabilities with high confidence except for the border pixels between adjacent targets (Fig 4C,E). The habitat maps from the DCRF-processed probability maps better delineated the benthic scene with smooth and contiguous regions (Fig 4A-B).

**Fig. 4.**
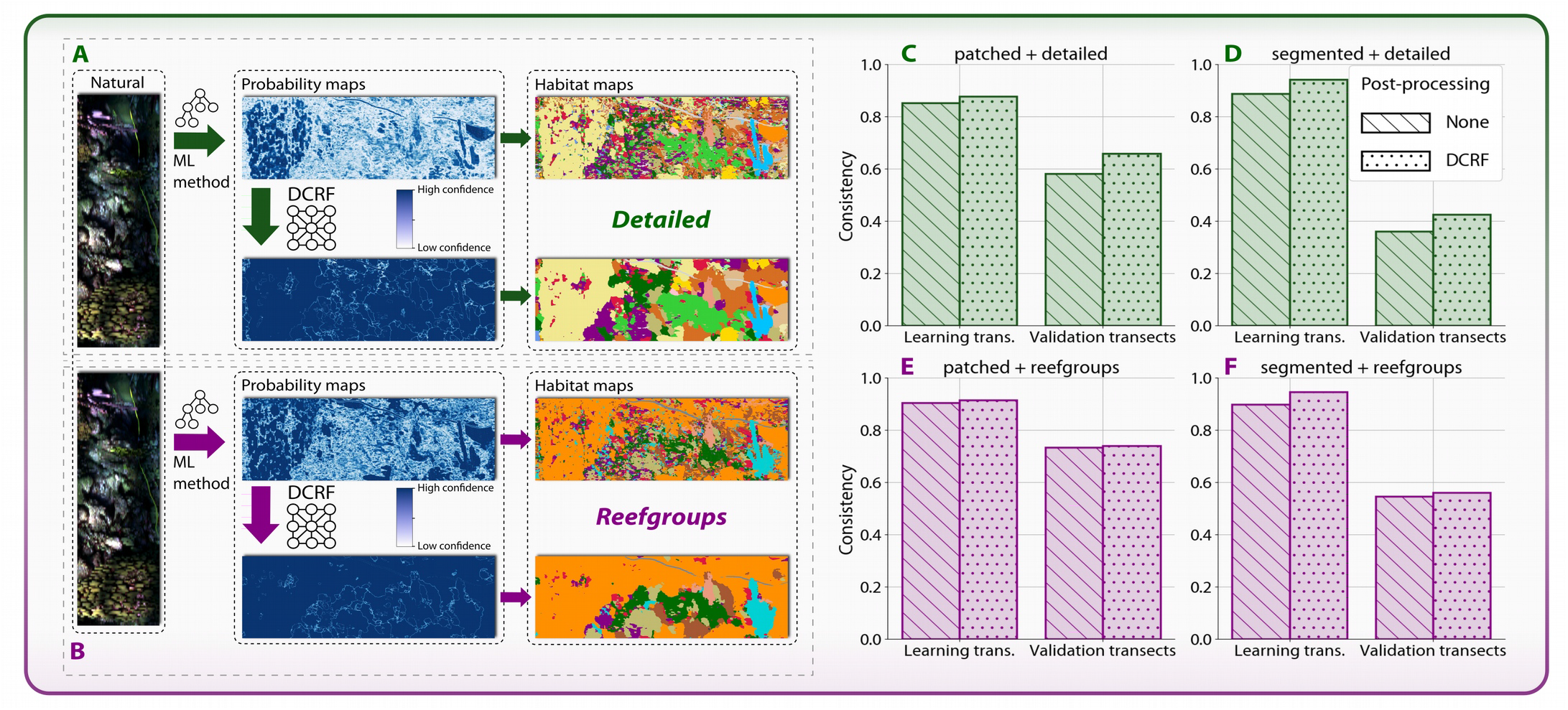
Smooth and consistent habitat mapping. The process of producing smoothed habitat maps in the detailed **(A)** and reefgroups **(B)** labelspaces. Habitat maps produced from the classifiers’ probability maps exhibit small-scale spatial noise and regions of low confidence. Smoothing the label probabilities with DCRFs renders spatially cohesive maps. **(C-F)** The use of DCRFs improved the map consistency in all cases. The patched method was better at generalization and was more accurate at mapping the validation transects, which were completely disjoint from the learning transects used for training the ML classifiers.

Beyond the visual cohesiveness, the smoothed habitat maps were quantitatively more consistent than the raw habitat maps in all combinations of ML methods and labelspaces for transects (Fig 4C,F; Fig S9A-B). The map consistency was measured as the average of the label accuracy in each of the annotation regions in all of the annotated transects, i.e. regions used for both training and testing the ML methods. The consistency of the habitat maps in the regions of the validation transects were lower than in the learning transects (Fig 4C-F): consistency for the segmented method dropped from 94% to 43% and from 95% to 56% with the detailed and reefgroups labelspaces (Fig 4D,F), and from 88% to 66% and from 92% to 74% for the patched method (Fig 4C,E). This indicated that the patched method (convolutional neural networks) was better than the segmented method (ensemble object classifiers) at generalizing to unseen data. Classification of transects with the reflectance signals resulted in more extreme drops in consistency up to 76% (Fig S9). Overall, the best predictive performance on data from the validation transects, which was not used in any ML step, was from the patched method.

Smooth habitat maps in both labelspaces were produced for all 31 transects. With each transect approximately 50 m × 1 m in size, this task involved assigning each of the 500+ million spectral pixels one of 43 labels (detailed) or one of 11 labels (reefgroups) independently. Montages of the habitat maps for all transects were visualized (Fig S10-S13). A selection of interesting sections of these habitat maps was collected (Fig 5) with the natural view (rows 1 and 4), the detailed map (rows 2 and 5) and the reefgroups map (rows 3 and 6) shown together. In image A:2 and A:3, the habitat maps show well-delineated instances of three coral species (*D. strigosa, P. asteroides* and *M. cavernosa*), a sponge (*N. nolitangere*) as well as regions of *Sediment, Turf algae* and *Dictyota* macroalgae. The maps in F:5 and F:6 show another example of fine-grained segmentation of the branches of a specimen of the *Plexauridae* soft coral family. Comparing the maps in D:2 and D:3, or in H:5 and H:6, shows the number of different coral species that can be identified under the broad *Coral* functional group (in orange). Small encrusting taxa such as *Coralline algae* are visible in B:2 and E:2. The delineation of *Cyanobacterial mat* in G:2 and I:2, along with the many regions of *Sediment* and *Turf algae* represent substrate and microbial components of coral reef benthos. The habitat map sections also display some confusions: different sections of the same colony in C:2 are assigned to *Plexauridae* and *Gorgoniidae,* which are both soft coral families with similar digitate morphologies, while in C:3 this same colony is assigned between *Coral* and *Soft coral.* The incorrectly labelled regions of *N. nolitangere* sponge in E:2 (and *Sponge* in E:3) along the image edge are also errors. These errors likely occur due to the poorly illuminated shadow regions that received predictions with low confidence, and then got reassigned to the *Sponge* class by the DCRF process due to nearby high-confidence regions. Note however that this did not occur for the *Transect tape* in the shadow regions which was predicted with high confidence.

**Fig. 5.**
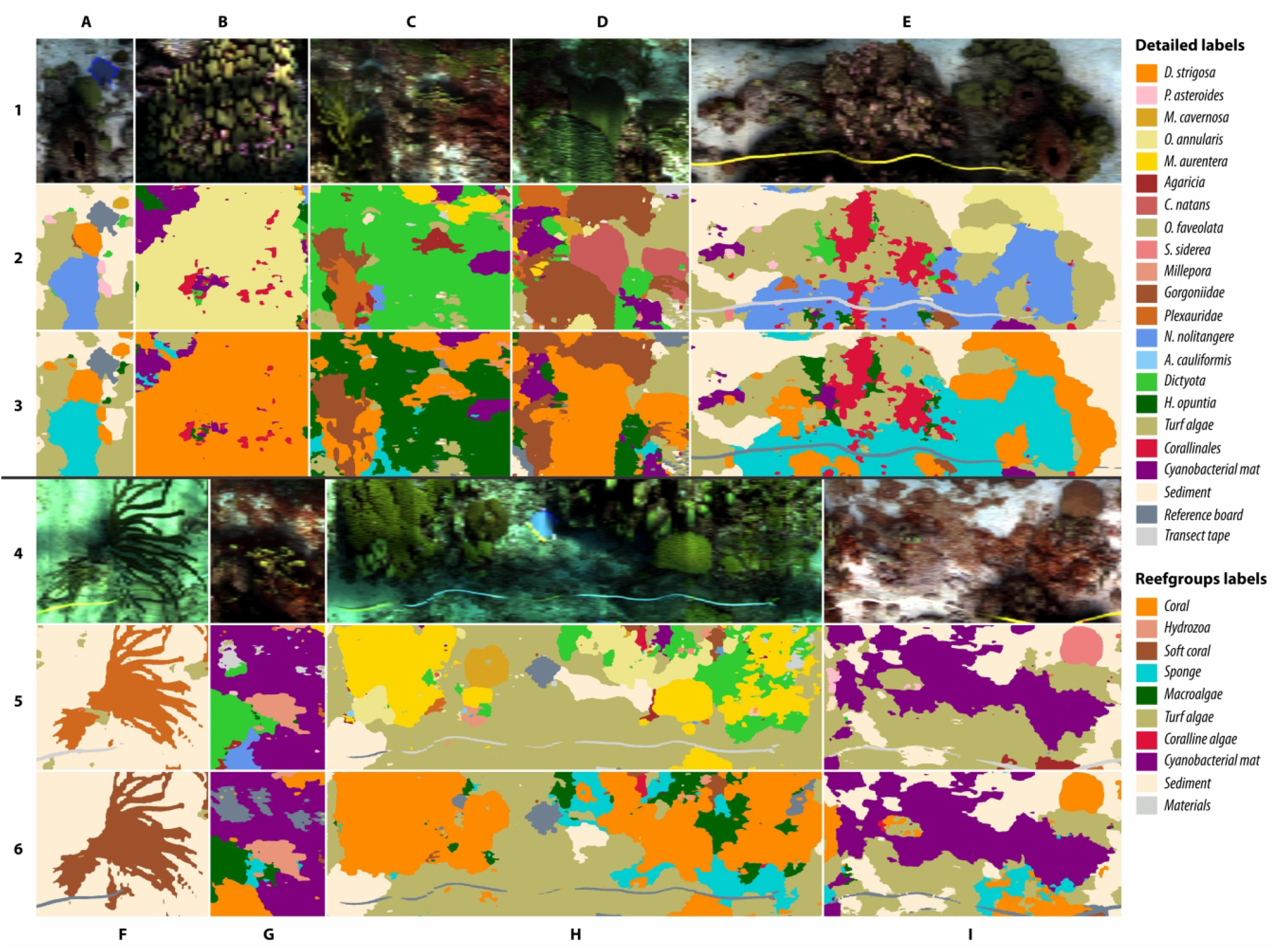
The rich structure of a digitized reefscape. Sections of habitat maps produced with the patched ML method, with colors corresponding to classes from each labelspace (see legend). Rows 1 and 4 are the natural view, as would be seen by a human observer. Rows 2 and 5 show the sections in the detailed labelspace and rows 3 and 6 in the reefgroups labelspace. Our proposed workflow accurately discerns among a large labelspace and delineates complicated shapes of reef biota. Detailed descriptions are in the *Results.*

### Assessing the habitat structure

A Bland-Altman analysis of the coral coverage (Fig 6A,B) and the Shannon diversity index (Fig 6C,D) from the reefgroups labelspace maps showed a high degree of correlation and low bias between the segmented and patched methods. The coverage of all other reefgroups classes, derived from either mapping method, were comparable across the range of values in all the transects (Fig S14). The Bray-Curtis similarity, which compares the compositional structure between the communities in the maps of both methods, had a median value of 82% in the detailed and reefgroups labelspaces with a quartile range of 72%-89% and 75%-90%, respectively (Fig 6E). With the *Sediment* and *Turf algae* substrate classes merged together, the median value for the similarity rose to 89% for the detailed and reefgroups labelspaces, with a quartile range of 78%-92% and 79%-94%, respectively. This improvement in the similarity index indicates a large effect of the inherent definition problem of *Turf algae* on reef habitat mapping (see *Discussion*).

**Fig. 6.**
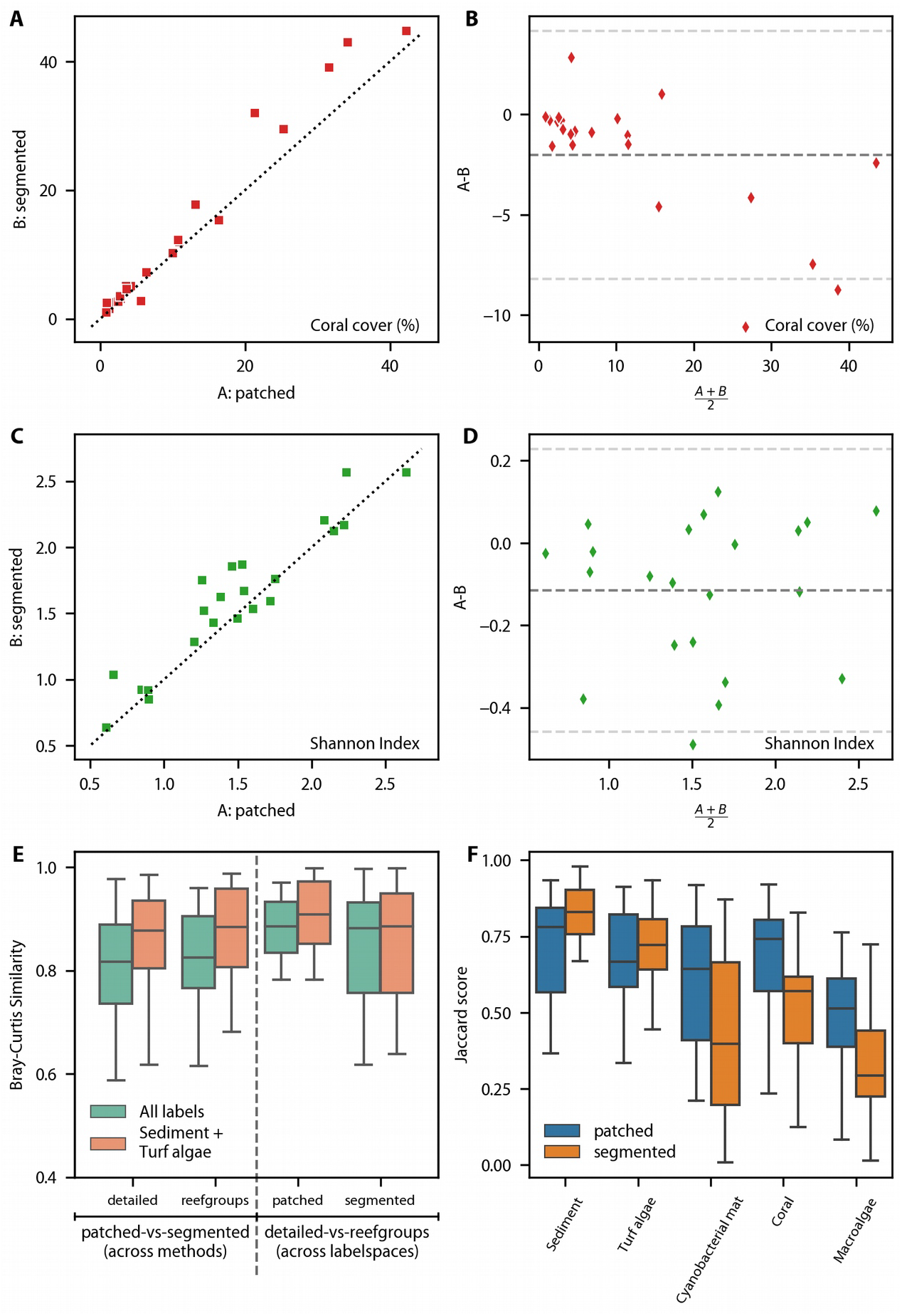
Assessing the benthic habitat structure from dense maps. **(A)** Coral cover of 22 learning transects was compared from the habitat maps for each ML method. The segmented method predicted slightly more coral cover than the patched method in the transects. **(B)** No clear bias was noted for either method. **(C)** The Shannon index values for 22 compared transects were highly correlated between the maps from the segmented and patched methods. The segmented method produced habitat maps with slightly higher Shannon diversity. **(D)** No significant bias in either ML method for the Shannon index comparison. **(E)** The median Bray-Curtis similarity between the mapped communities was ~80% across ML methods was and ~88% across labelspaces. Note that the orange bars refer to a labelspace where *Sediment* and *Turf algae* were combined into a single class, resulting in a higher compositional similarity. **(F)** Configurational similarity assessed using the Jaccard index between the reefgroups and detailed-to-reefgroups habitat maps for the top-five dominant labels are shown. Given the 500+ millions of pixels in this assessment, the maps showed very good consistency in configuration.

The similarity assessment across the learning transects for the patched method showed an 88% similarity median value with a quartile range of 84%-92%, while the segmented method showed an 87% median value with a quartile range of 77-91%. With the *Sediment* and *Turf algae* classes merged, the patched method showed a similarity median of 90% (quartile range 88%-97%), whereas the segmented method showed a barely improved median similarity of 88% (quartile range 77%-93%). Therefore, our proposed workflow recovered a reef community composition that was highly consistent between the taxonomic and functional group descriptions of the reef benthos.

The spatial configuration analysis (Fig 6F) between the detailed-to-reefgroups and reefgroups maps showed that three classes had high configurational similarity for the patched method, *Cyanobacterial mat* (64% vs. 40%), *Coral* (74% vs. 57%) and *Macroalgae* (51% vs. 29%). Two classes showed better configurational similarity for the segmented method, *Sediment* (83% vs. 78%) and *Turf algae* (72% vs. 67%). The Jaccard score was lower for both methods on rarer classes (Fig S15).

### Evaluating the effort-vs-error compromise in reef sampling

We exploited our dense and accurate habitat maps to revisit the effort-vs-error relationship of sparser reef sampling techniques. The number of point samples required to achieve a relative error lesser than 5% was assessed (Fig 7). To recover the hard coral coverage in transect T1 – which had low biodiversity (H’ = 0.6) and low coral coverage (2.3%) – 960 points per quadrat were required (Fig 7B,C). For the transect T4, with H’ = 2.6 and coral coverage of 34.2%, 80 points per quadrat were sufficient. Similarly, 960 points per quadrat were needed to recover the sediment coverage in transect T4, where sediment covers only 2% of the benthos (Fig 7D,E). In contrast, only 5 points per quadrat were required to capture the sediment coverage in transect T1, which has the highest sediment coverage (84.6%). The Shannon index was recovered with 10 points per quadrat for transect T4 (H’ = 2.6), with 40 points per quadrat for the transects T2 (H’ = 1.6) and T3 (H’ = 1.3), and with 160 points per quadrat for transect T1 (H’ = 0.6) (Fig 7F,G). Overall, higher sampling effort was required to accurately recover the coverage of rare species or to capture the Shannon biodiversity index of scenes with low biotic coverage.

**Fig. 7.**
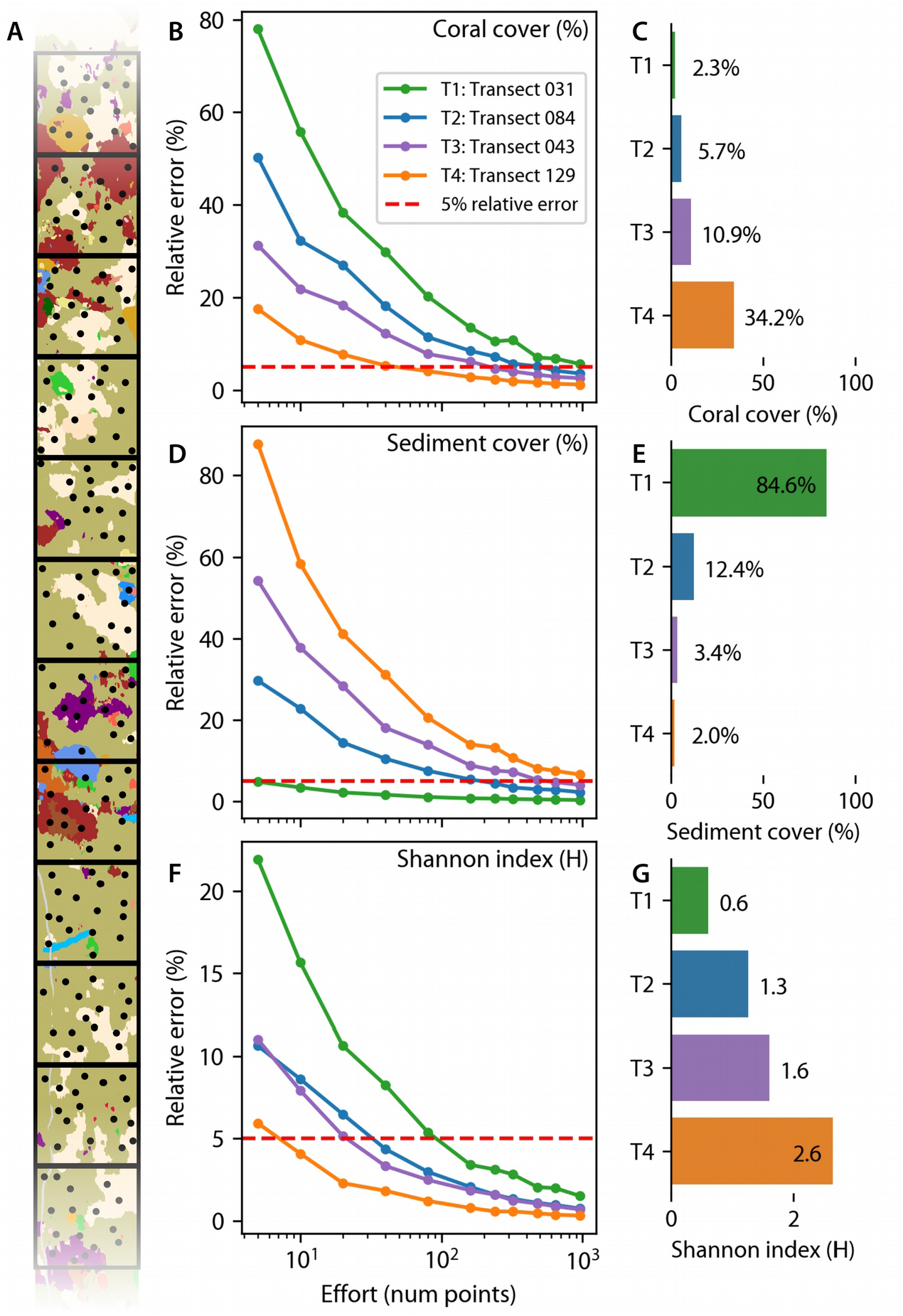
Effort-vs-error analysis for point-count sampling of reef habitat structure. **(A)** Schematic of the simulations simulations of quadrat-wise random sparse sampling of dense habitat maps to estimate metrics (coverage or Shannon index) through point-count estimates of four transects with different biodiversity values (H = {0.61, 1.26, 1.6, 2.64}). The error in the habitat metric from sparse random points relative to the metric of the full map was calculated from repeated trials. The number of point samples per quadrat required to achieve a relative error lesser than 5% (dashed line) was assessed. **(B, C)** Over 960 points to estimate coral coverage in T1. **(D, E)** Over 960 points to estimate sediment coverage in T4. **(F, G)** Over 80 points are required to correctly estimate Shannon index in T1. These results suggest that rarer species require more sampling effort and that over 80 points per quadrat should be used to estimate habitat metrics in reef transects where the expected diversity or coverage is not previously known.

## Discussion

The presented reef mapping workflow was able to produce dense habitat maps with an unprecedented degree of thematic (43 labels) and spatial (~2.5 cm/pixel) detail. The rich spectral detail in the HyperDiver data was leveraged by machine learning classifiers to produce highly accurate habitat maps (87% Fbeta) with little annotation effort (2% pixels). The two labelspaces used in the maps describe the reef benthic biodiversity down to genus and species level as well as abiotic and microbial components, such as turf algae and cyanobacterial mats. Our habitat maps provide a no-pixel-left-behind dense view of entire 50 m long transects, which allowed us to localize and identify the visible components of the reef benthic community. Two ML methods with different operational paradigms were used independently to produce highly accurate habitat maps, thus enabling an objective comparison of reef descriptions at big data scale. Our workflow generated a deep description of habitat structure (diversity, coverage, composition and configuration), which demonstrated high convergence between both ML methods. Nonetheless, our detailed assessment indicates that deep learning classifiers are better at generalizing towards new and unseen datasets under comparable annotation and computational effort.

Our workflow uses a detailed thematic scale in the form of 43 benthic categories that describe biotic and abiotic components of the reef habitat. Subsequently, a reefgroups labelspace, comprised of functional groups of reef biodiversity, was abstracted from the taxonomically detailed labelspace through interconnected ontologies, similar to a hierarchical geomorphic zones developed previously (Kennedy et al., 2021). By independently mapping into the reefgroups labelspace we showed that the workflow consistently retrieved the composition and configuration of the reef transects across thematic resolutions (Fig 6E,F). This allows for the workflow to ‘translate’ between the needs of different expert groups, such as reef ecologists or managers (Lecours et al., 2015; Roelfsema et al., 2021).

To automate the digitization of reef structure, careful consideration of workflow parameters is recommended. In our workflow, we independently utilized two ML approaches: one based on deep neural networks and the other on object-based image analysis. Continual increase in complexity and specificity of ML tasks for automation impedes a clear judgment of an ML method’s ability to generalize (Paoletti et al., 2019). Following best practices, we tested two ML methods with different operational paradigms on a non-overlapping subset of learning data to explore the trade-off in performance vs. complexity of automation. Both segmented and patched methods showed 80% ± 5% weighted accuracies in both labelspaces (Fig 2), the segmented approach performing better with reflectance data and the patched method with radiance data. Another work targeting a similar sized labelspace (35 labels) achieves a mean pixel accuracy of 49.9% with a semantic segmentation network (DeepLabV3+) on sparse samples in 729 test images of coral reef orthomosaics (Alonso et al., 2019). Our workflow, given only 2% of annotated pixels and underwater transects with high variability in imaging settings, produced an impressive >75% accuracy in classifying 43 labels. We show that relatively simple and complex ML methods can produce accurate mapping of reef transects, apparently due to the spectral detail (Fig 3C).

Another methodological focus was the data requirements for the ML algorithms. We assessed the performance of both methods – under a constant computational effort – based on the number of unique pixels (in segments or patches) used for training (Fig 3A,B). The patched method needed less data to achieve its peak performance under the same amount of computing power and annotation effort. Despite the variable lighting conditions and methodological artefacts between training and validation transects, the patched method classified into both labelspaces more consistently (Fig 4C-F). Although classifier-level metrics are better for the segmented method (Fig 2), the patched method was 23% better at classifying out-of-distribution data in the detailed labelspace and 18% better in the reefgroups labelspace than the segmented method (Fig 4C-D). Given that expert annotation is the biggest bottleneck for reef survey analysis (Beijbom et al., 2015; Roelfsema & Phinn, 2013), the patched method provides better performance-per-human-effort compared to the segmented method.

The smoothed habitat maps from our workflow show spatial and thematic detail of the structure of the coral reef benthic community (Fig 5). Benthic targets are clearly separated into meaningful regions, which represent different substrata, different organisms of various sizes and different taxonomic classes. Small coral colonies and intricate shapes of branching corals, soft corals and sponges, and even transect tapes are correctly delineated and classified (see *Results*). Our workflow was able to accurately map bare sediment, turf algae and cyanobacterial mats achieving a previously missing capability in reef habitat mapping: dense mapping of the microbial components of reef substrata, while integrating them into a benthic community labelspace. This can be used to quantify changes in abundance of cyanobacterial mats or turf algae, usually an indicator of reef deterioration (de Bakker et al., 2017). Our workflow achieved a dense no-pixel-left-behind mapping of the intricate spatial structure and taxonomic diversity of the micro- and macro-benthic communities in coral reefs.

As the primary focus of our workflow development was to convert underwater spectral images into dense habitat maps, the target of our assessment was to go beyond comparing classifier-level metrics, and assess the final habitat maps for consistency and accuracy. We considered the goal of comparing the densely classified maps to photo-quadrats with point-count estimates (Rashid & Chennu, 2020), but the large difference in sample sizes – ~2000 quadrat points vs. millions of classified points – would not be statistically sound. Given that 98% pixels (out of 500+ million) do not have reference label annotations and the complex spatial structure of the transects (Fig S10-S13), it is difficult to assess the maps accuracy on a pixel level. To overcome this limitation, we exploited the fact that the two ML methods produced the same type of output, but worked fully independent of each other in terms of method, input signal and parametrization. We compared the habitat structure of the dense maps in describing the same physical transect of the seafloor. To achieve this we used coverage, as well as composition and configuration metrics, which are key descriptors of habitat structure (Nowosad & Stepinski, 2019; Riitters, 2019). We consider that if the statistical properties of the habitat maps from independent methods are similar, our workflow will have succeeded in representing the true composition and configuration of the coral reef transects captured by the underwater surveys.

The Bland-Altman analysis showed that the habitat maps agree on coverage and diversity metrics, except for some small discrepancies (Fig 6A,B; Fig S14). Coverage of corals, sponges and cyanobacterial mats were overestimated by the segmented method in some transects (Fig S14 I-J). The composition of the communities between pairs of dense habitat maps (considering all pixels) was highly similar as shown by the Bray-Curtis similarity index (Fig 6E). This shows that both ML methods independently ascribe similar classes and similar number of those classes to the same transect. Furthermore, the assessment of the communities between the two labelspaces was also very similar (Fig 6E), providing confidence in the thematic flexibility of our workflow. We infer that automated mapping with ML methods of underwater hyperspectral transects can handle intra-class variability (detailed labelspace) as well as inter-group variability (reefgroups labelspace) with high accuracy.

Going beyond the composition, we also assessed the spatial configuration similarity of the benthic community in the transects described by both ML methods using the Jaccard index. The regions from the abstracted detailed-to-reefgroups maps and direct reefgroups maps yielded Jaccard scores over 60% for the dominant labels (Fig 6F). Given that these comparisons are across hundreds of millions of pixels and over 43 different labels, these results indicate high configurational similarity between the maps. Therefore, our workflow is able to correctly localize and delineate important targets in benthic habitat maps, despite the degree of thematic detail. Nonetheless, these assessments are inter-comparisons within our workflow and a correct assessment of the configuration of the habitat structure would require dense manual annotations of the transects. The high degree of convergence in the habitat structure mapped independently by the two ML methods using two different hyperspectral signal types gives confidence in the ability of the workflow to produce accurate in-situ descriptions of coral reef habitats with unprecedented detail and analytical throughput.

There has been some debate about the amount of sampling effort necessary to accurately measure reef habitat structure. In benthic surveys with photo-quadrats and point count analysis, the ‘adequate’ number of samples (points) per quadrat to accurately describe the community has been a topic of debate (Dumas et al., 2009; Pante & Dustan, 2012; Perkins, 2016). The goal is to find a good balance between expert effort (labelling the points in images) and reliability in the derived scene description. Different numbers (from 5 points to 99 points per quadrat) have been proposed and used in survey analyses (Dumas et al., 2009). Simulations of sampling synthetic habitat maps, based on normal distribution of class abundances, to recover the coral and sponge coverage estimates with less than 5% relative error, revealed that the optimal number of points per quadrat ranged between 13 points for a heterogeneous area with high coral coverage and over 600 points per quadrat for a very homogeneous region with low coral coverage (Pante & Dustan, 2012). The recommended number of points per quadrat was 80 for transects of unknown community structure, but generally depended on the true underlying diversity and dispersion of the community configuration.

We contribute to the debate with a reef-scale analysis based on empirical community structure derived from our benthic habitat maps (Fig 7). We simulated quadrat sampling with various degrees of sampling effort, and found that the number of point per quadrat to accurately recover the coverage of rare species exceeded the recommended values in the literature. Even with 1000 sampling points per quadrat, rare species at the transect level could not be detected within the 5% relative error limit (Fig 7B,D). In contrast to the coverage of individual classes, the Shannon diversity index was captured within 5% relative error with 160 sampling points per quadrat (Fig 7E). We demonstrate that the rarity and skewness of occurrence significantly impacts the error associated with a constant sampling effort. When dense mapping is not possible, and no prior knowledge of the biodiversity at a site exists, we recommend to sample over 160 points per quadrat during generation of baseline data, as well as to communicate the uncertainty generated from this sampling design (Hochberg & Gierach, 2021). If rare species are an important survey target, then dense habitat mapping represents the least effort towards accurate estimations of species coverage. Finally, dense mapping allows to describe species distributions in more detail and goes beyond reef functional groups demographics, which can hide intra-group shifts and conceal key dynamics of coral reef communities (Brito-Millán et al., 2019).

## Conclusions

Our proposed workflow showcases a way to generate dense habitat maps of coral reefs with flexible thematic detail. This thematic and spatial detail in the maps enables fine-grained analyses of coral reef functions and community dynamics by coral reef ecologists. We seek to unify the perspective of ecologists, environmental managers, remote sensing and machine learning communities involved in the study of coral reefs. Particularly for ecologists and managers, it provides a consistent habitat description with adaptive thematic detail. Between remote sensing and machine learning experts, it offers a perspective on bridging the ‘measurement gap’ between ML classifiers and the ultimate data products, i.e. habitat maps. The consistency achieved by our mapping workflow, and the patched method in particular, is related to the richness in spectral detail and the spatial acuity of our proximal sensing vantage point of underwater surveys (Chennu et al., 2017; Rashid & Chennu, 2020). When certain limitations are overcome (see *Supplement*) and with improvements in cost and performance of underwater spectral surveying technology, it will become feasible to integrate it as a standard in-situ reef monitoring technique. The widespread use of underwater spectral surveying and automated benthic habitat mapping promises to provide the best validation data for aerial Earth observation efforts to map coral reefs globally. The integration of thematic detail into global habitat mapping promises to enable novel analyses of pattern and scale in coral reef ecology.

## Acknowledgments

We would like to thank Carsten John and Oliver Artmann at the Max Planck Institute for Marine Microbiology for their IT support. This project has received funding from the European Union’s Horizon 2020 research and innovation programme under the Marie Sklodowska-Curie grant agreement number 813360 “4D-REEF”.

## Data availability statement

All data needed to evaluate the conclusions in the paper are present in the paper, the Supplementary Materials and in a public data repository (undergoing submission at the Pangaea repository).

## Author contributions

Daniel Schürholz – Methodology, Software, Validation, Formal analysis, Investigation, Data curation, Writing – Original Draft, Writing – Review & Editing, Visualization, Project administration.

Arjun Chennu – Conceptualization, Methodology, Software, Validation, Formal analysis, Investigation, Data curation, Writing – Original Draft, Writing – Review & Editing, Visualization, Supervision, Project administration, Funding acquisition.

