## Supplemental text, figures and tables. for "Digitizing the coral reef: machine learning of underwater spectral images enables dense taxonomic mapping of benthic habitats"


**This PDF file includes:**

Spectral-spatial neural network parameters and machine learning libraries
Limitations of our workflow
Table S1
Figures S1 to S15
Supplement references

### Spectral-spatial neural network parameters and machine learning libraries

The parameters used for training the spectral-spatial neural network network were: a batch size of 32, a number of epochs of 32 and an input patch size of 13×13 pixels. Cross entropy loss with weight balancing based on class abundance was used as the loss function during network training. The optimizer was a rectified Adam function with a weight decay value of 1.5 and had a β of 0.999 (Liu et al., 2021). The learning rate value was adjusted by a cyclic learning rate scheduler, oscillating from $1e^{-8}$ to $1e^{-3}$ in triangular ramps with a step of 250 batches of data samples. The whole network training workflow was built with the PyTorch library (Paszke et al., 2019), the Skorch machine learning workflow management library (https://github.com/skorch-dev/skorch) and the Mlflow training visualization tool (https://github.com/mlflow/mlflow/). Training was performed on a machine with two Nvidia RTX2080 GPUs, each with 12Gb dedicated memory.

The principal component analysis done for the segmented method was implemented using the scikit-learn library (Pedregosa et al., 2011). Furthermore, the watershed algorithm from the scikit-image library was used (van der Walt et al., 2014). Dense conditional random fields (DCRFS were implemented with the pyDenseCRF python library (https://github.com/lucasb-eyer/pydensecrf).

### Limitations of our workflow

In the main manuscript we presented an end-to-end workflow that produced consistent habitat maps for a large area of coral reefs. Although the results are encouraging enough to recommend the different methods applied in this work, some limitations are noteworthy. For example, the nature of the images gathered by the push-broom hyperspectral camera (without any georeferencing) meant that the resulting transects are not ideal for photogrammetric techniques, thus hindering the generation of orthomosaics and 3D models. Recent studies are investigating novel techniques to overcome this limitation and have succeeded in producing rectified orthomosaics of hyperspectral transects (Jurado et al., 2021; Moroni et al., 2012). With the blistering pace of development in machine learning, novel architectures and paradigms could be implemented to render better spatial-spectral classification with fewer annotations. Furthermore, image processing techniques that exploit underwater image formation models to account for the water column properties can aid standardization of data for ML (Akkaynak & Treibitz, 2019). Regardless of the data and ML method developments used in reef surveys, we consider it important to evaluate the dense habitat maps (and not the classifiers) in terms of accuracy and completeness, as the measure of progress. To disentangle the effects of changes in ML methods and data, we urge that the original images and annotations be made publicly available so that they can be re-evaluated independently.

### Tables

| Method | Labelspace | Signal type | Overall accuracy | Precision | Recall | Fbeta | Cohen's Kappa |
|---|---|---|---|---|---|---|---|
| *patched* | *detailed* | *radiance* | 0.767 | 0.779 | 0.769 | 0.769 | 0.760 |
| | | *reflectance* | 0.715 | 0.723 | 0.720 | 0.720 | 0.708 |
| | *reefgroups* | *radiance* | 0.869 | 0.871 | 0.869 | 0.869 | 0.829 |
| | | *reflectance* | 0.841 | 0.841 | 0.841 | 0.841 | 0.793 |
| *segmented* | *detailed* | *radiance* | 0.772 | 0.779 | 0.771 | 0.771 | 0.755 |
| | | *reflectance* | **0.846** | **0.848** | **0.845** | **0.845** | **0.835** |
| | *reefgroups* | *radiance* | 0.792 | 0.796 | 0.794 | 0.794 | 0.737 |
| | | *reflectance* | **0.875** | **0.876** | **0.875** | **0.875** | **0.841** |

**Table S1. Classifiers performance.**
Comparison of the performance of each ML method in combination with each labelspace and each signal type. The classifiers were tested on disjoint datasets of 50000 patches for the patched method and 15946 segments for the segmented method. The best performing classifiers are highlighted in boldface for the detailed and reefgroups labelspace.

### Figures

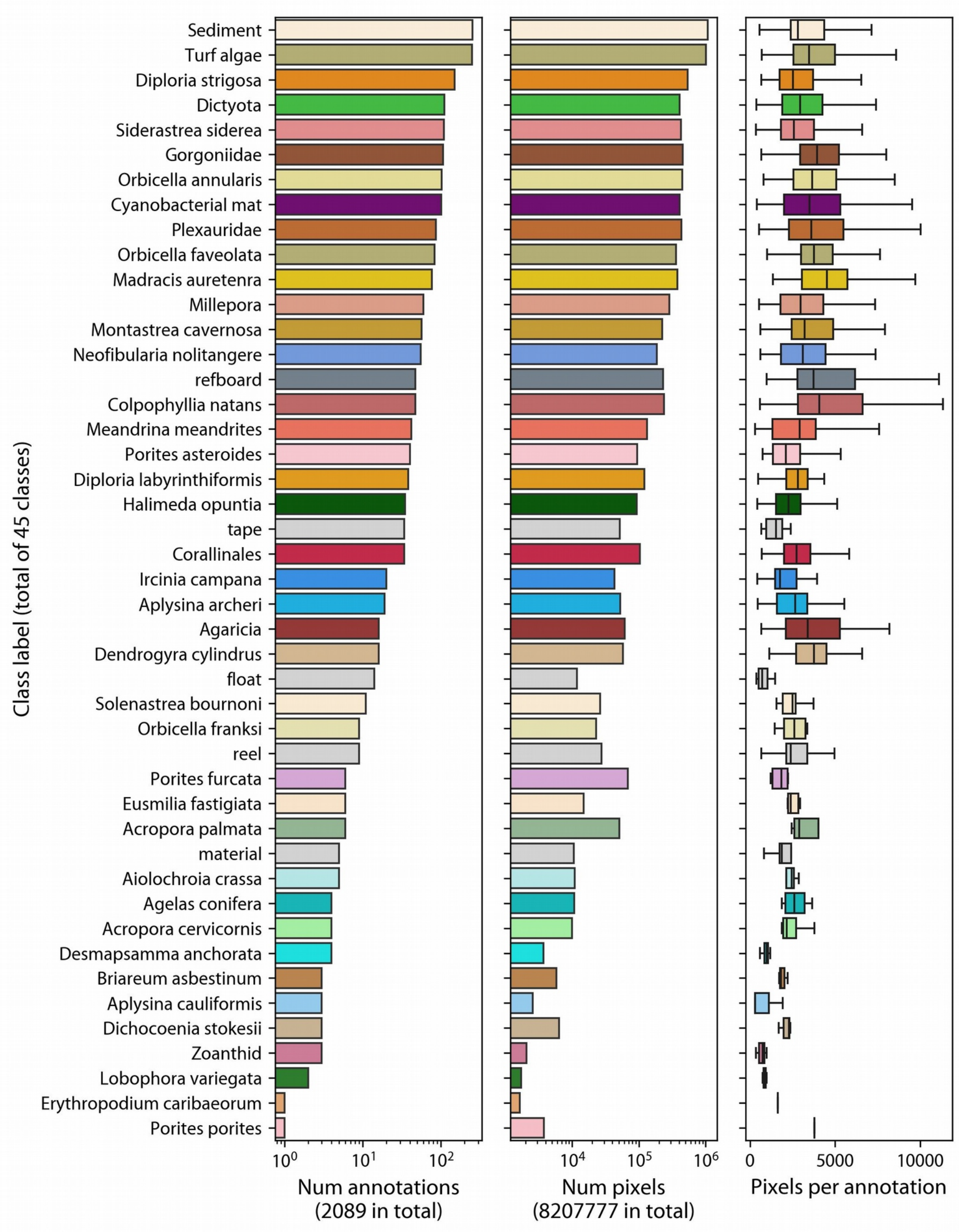


**Fig. S1.** Class distributions of detailed annotations. The dataset shows a large class imbalance in annotation regions in learning and validation transects. Classes such as *Sediment* and *Turf algae* are abundant in almost every transect with over 200 regions and covering around one million pixels. Rare classes such as *L. variegata* and *Zoanthid* have only 2 and 3 regions respectively, covering only around 2000 pixels. The annotations across classes were of similar sizes in term of number of pixels contained, except for rare species.

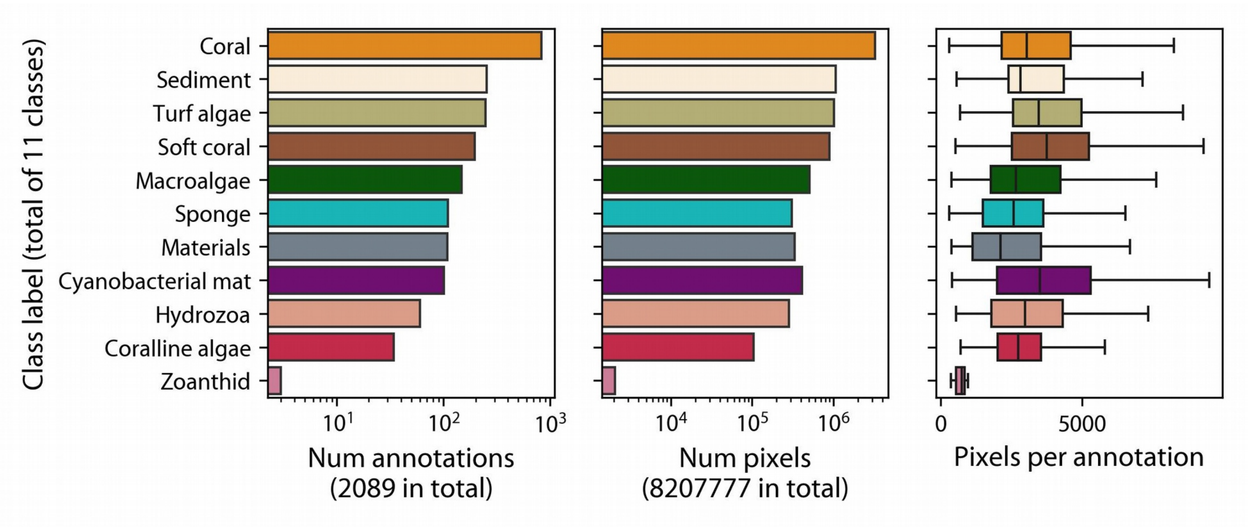


**Fig. S2.** Class distributions of reefgroups annotations. The reefgroups dataset less class imbalance than the detailed labelspace learning and validation transects. Classes such as *Coral*, *Sediment* and *Turf algae* are abundant in almost every transect with over 200 regions and covering around one million pixels. Rare classes such as *Zoanthid* have only 2 and 3 regions respectively, covering only around 2000 pixels. The annotations across classes were of similar sizes in term of number of pixels contained.

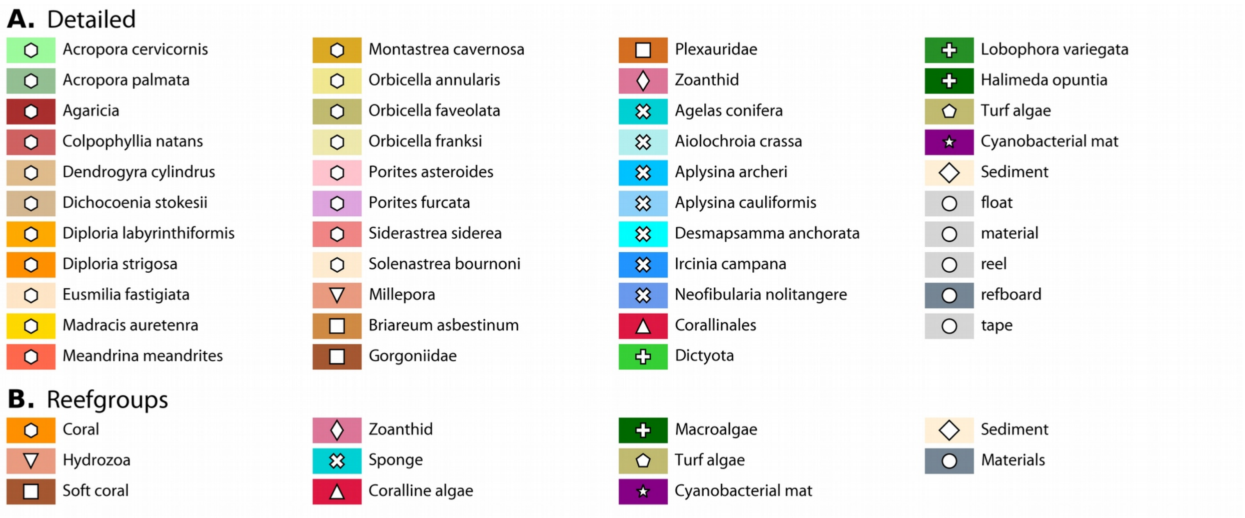


**Fig. S3.** Detailed and reefgroups labelspaces. **(A)** Reefgroups labelspace consisting of 11 labels for reef functional groups and abiotic elements. Some classes are divided into two or more classes in the detailed labelspace. Shapes within the color boxes are provided for easier identification between labelspaces. **(B)** Detailed labelspace consisting of 43 classes, representing the deepest possible taxonomic definition of targets in visual annotation of underwater transects.

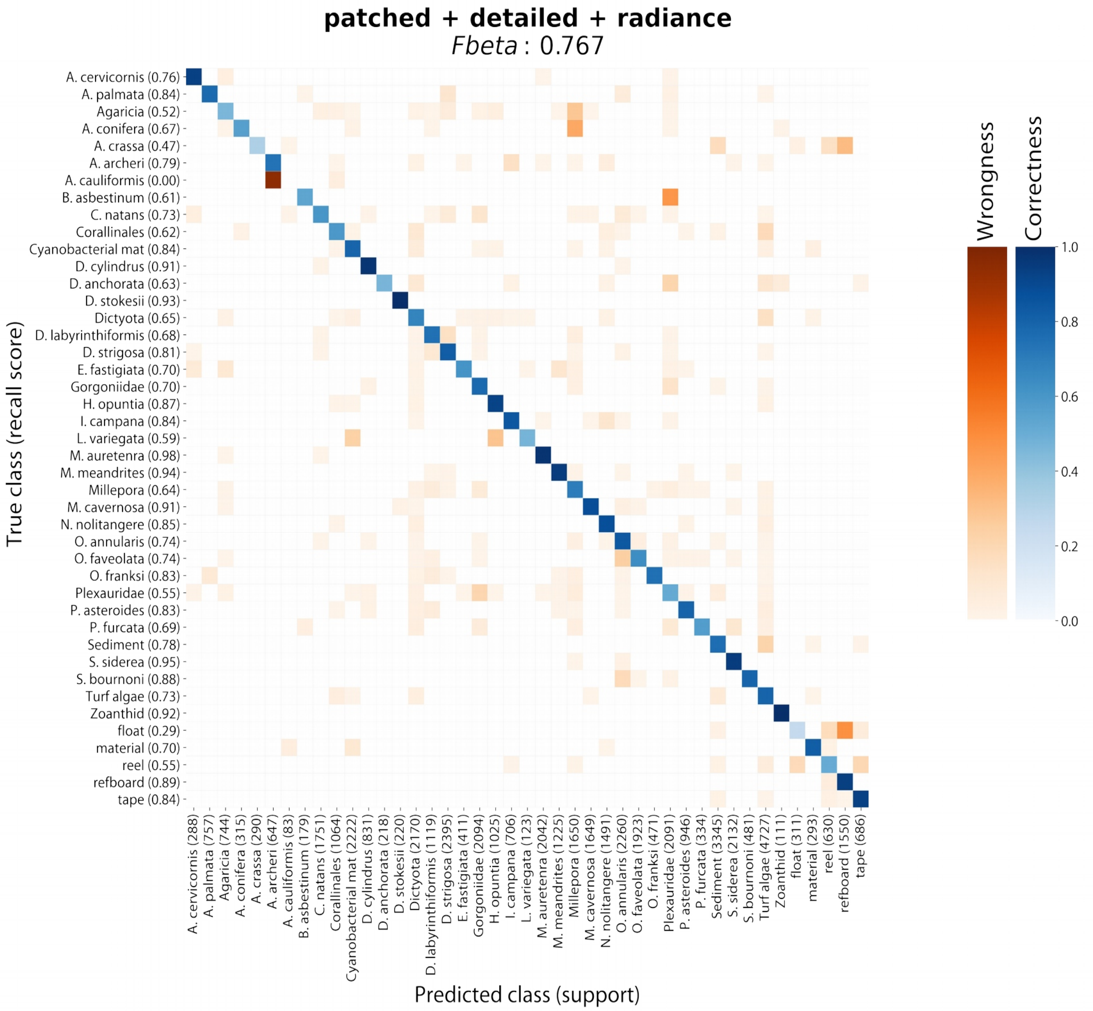


**Fig. S4.** Recall confusion matrix for patched classifier predicting into the detailed labelspace from radiance images.

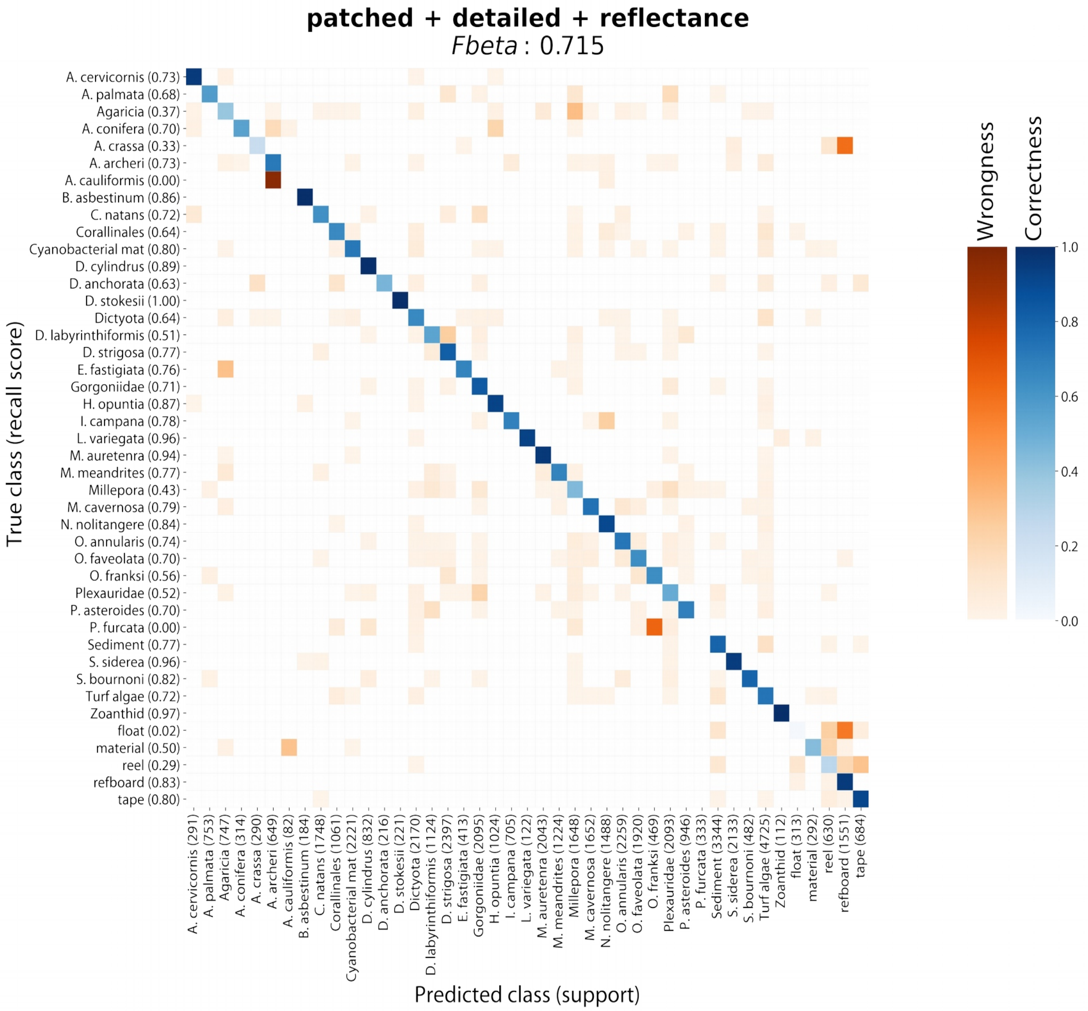


**Fig. S5.** Recall confusion matrix for patched classifier predicting into the detailed labelspace from reflectance images.

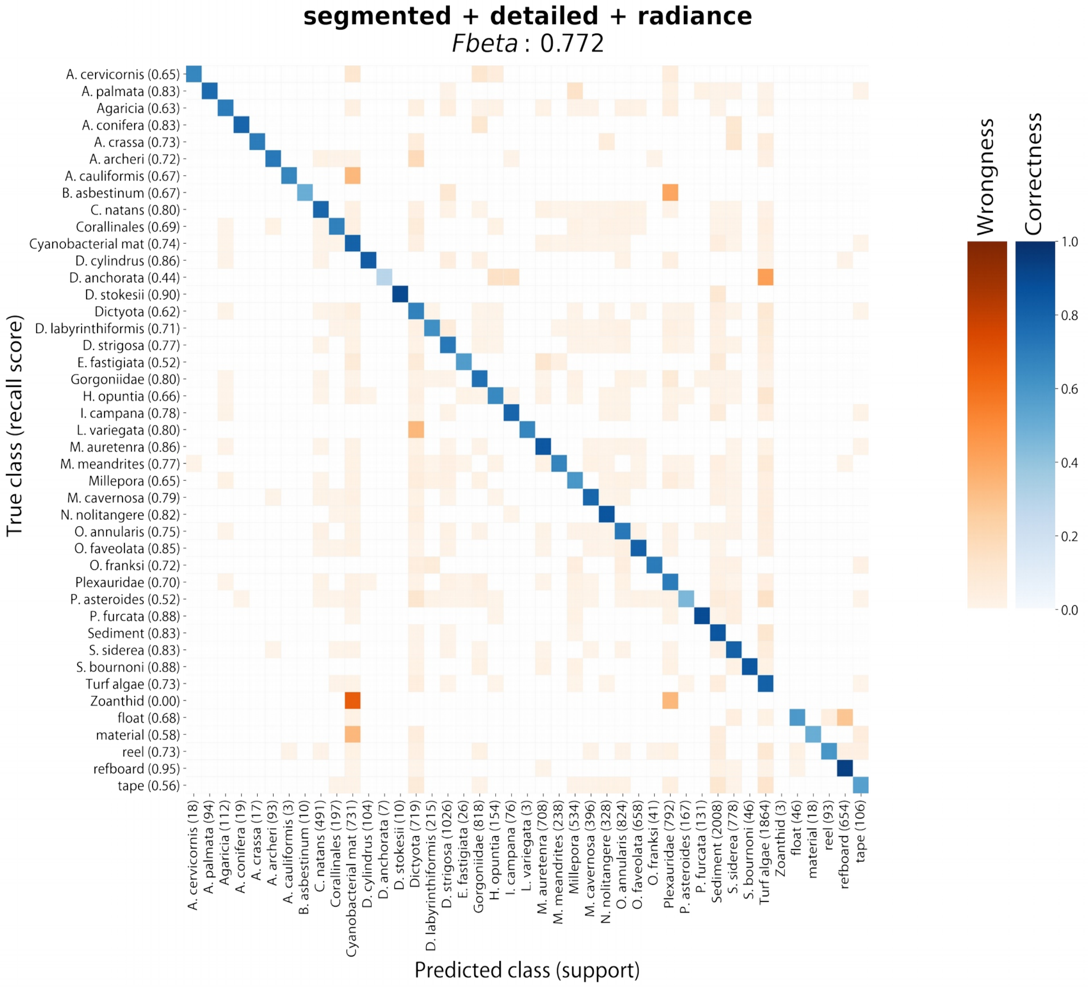


**Fig. S6.** Recall confusion matrix for segmented classifier predicting into the detailed labelspace from radiance images.

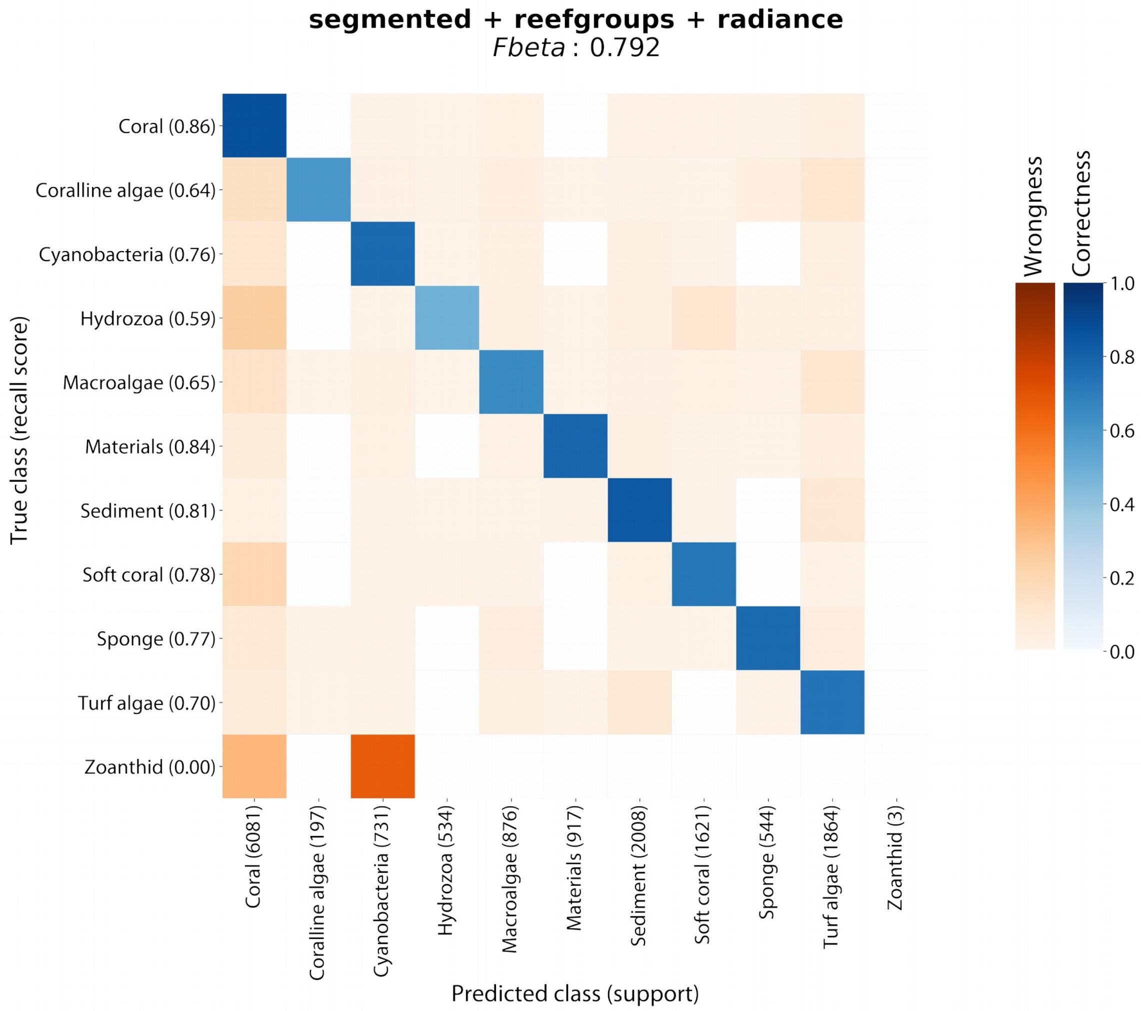


**Fig. S7.** Recall confusion matrix for segmented classifier predicting into the reefgroups labelspace from radiance images.

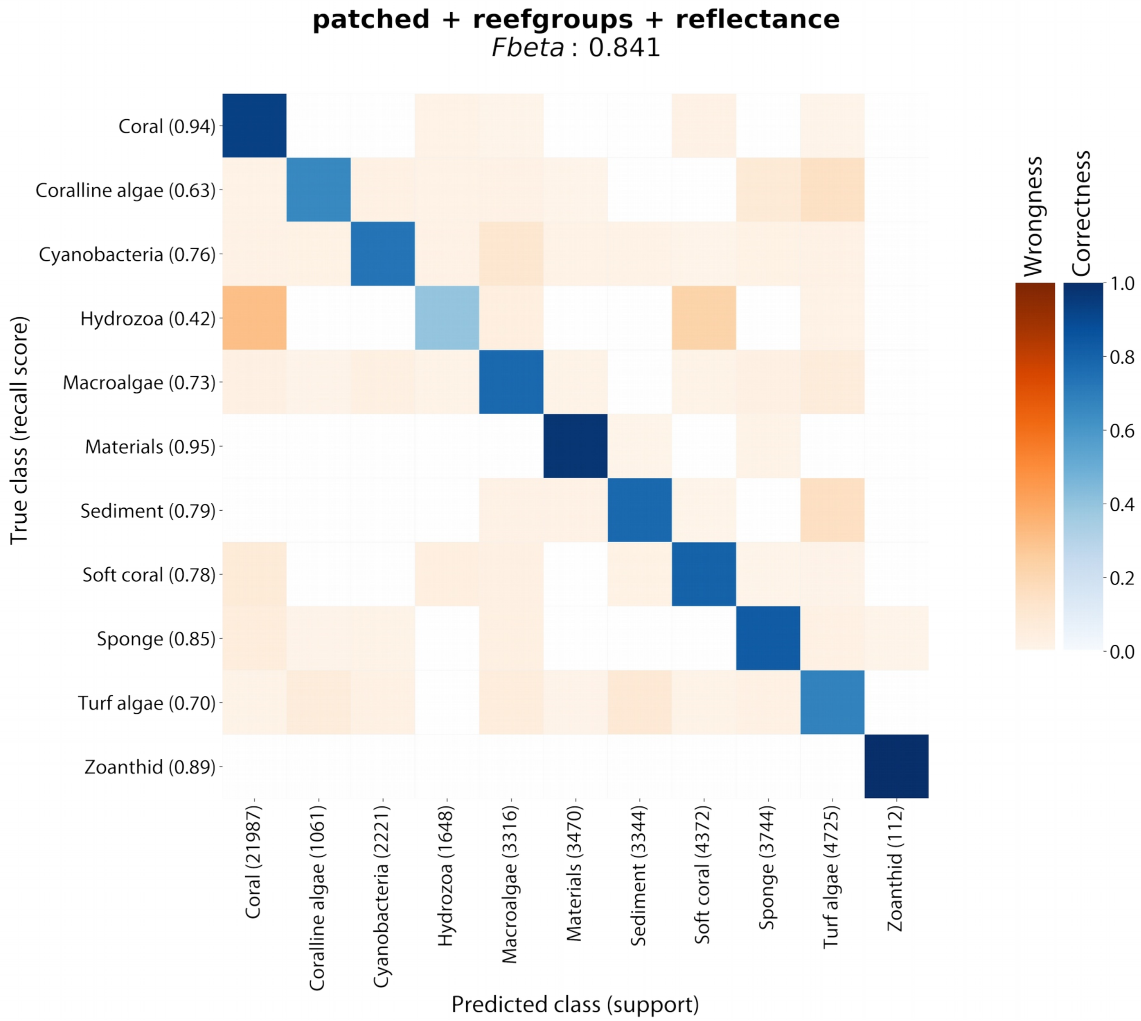


**Fig. S8.** Recall confusion matrix for patched classifier predicting into the reefgroups labelspace from reflectance images.

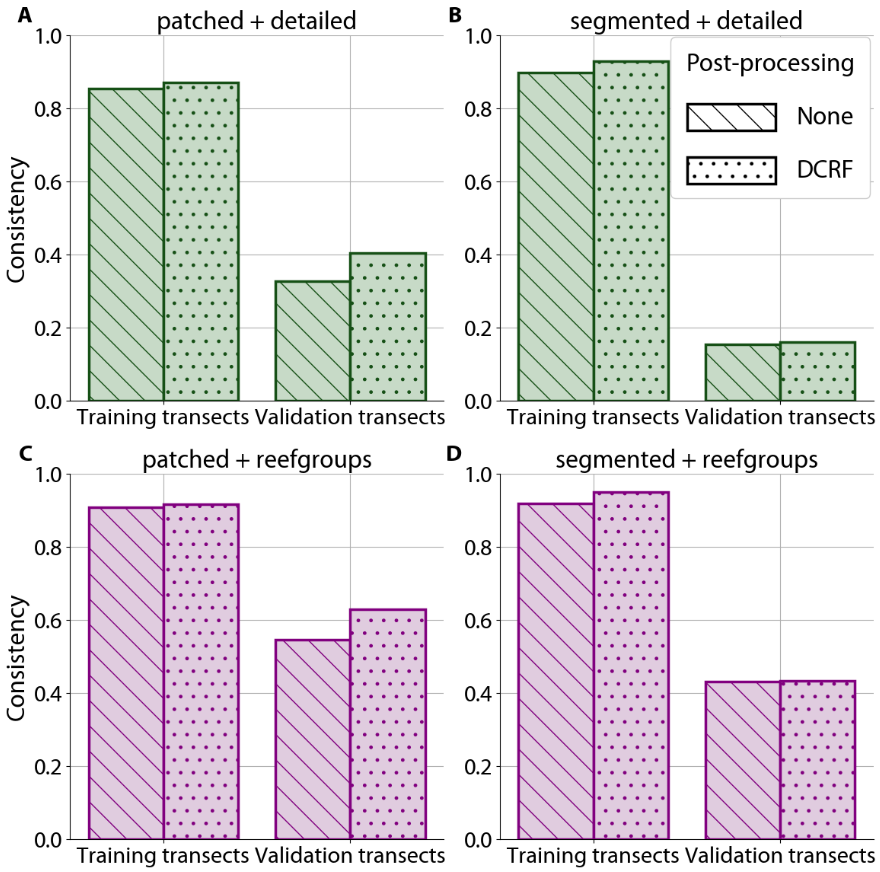


**Fig. S9.** Classification consistency for combinations of ML methods and labelspaces as in Fig. 4, but using reflectance data.

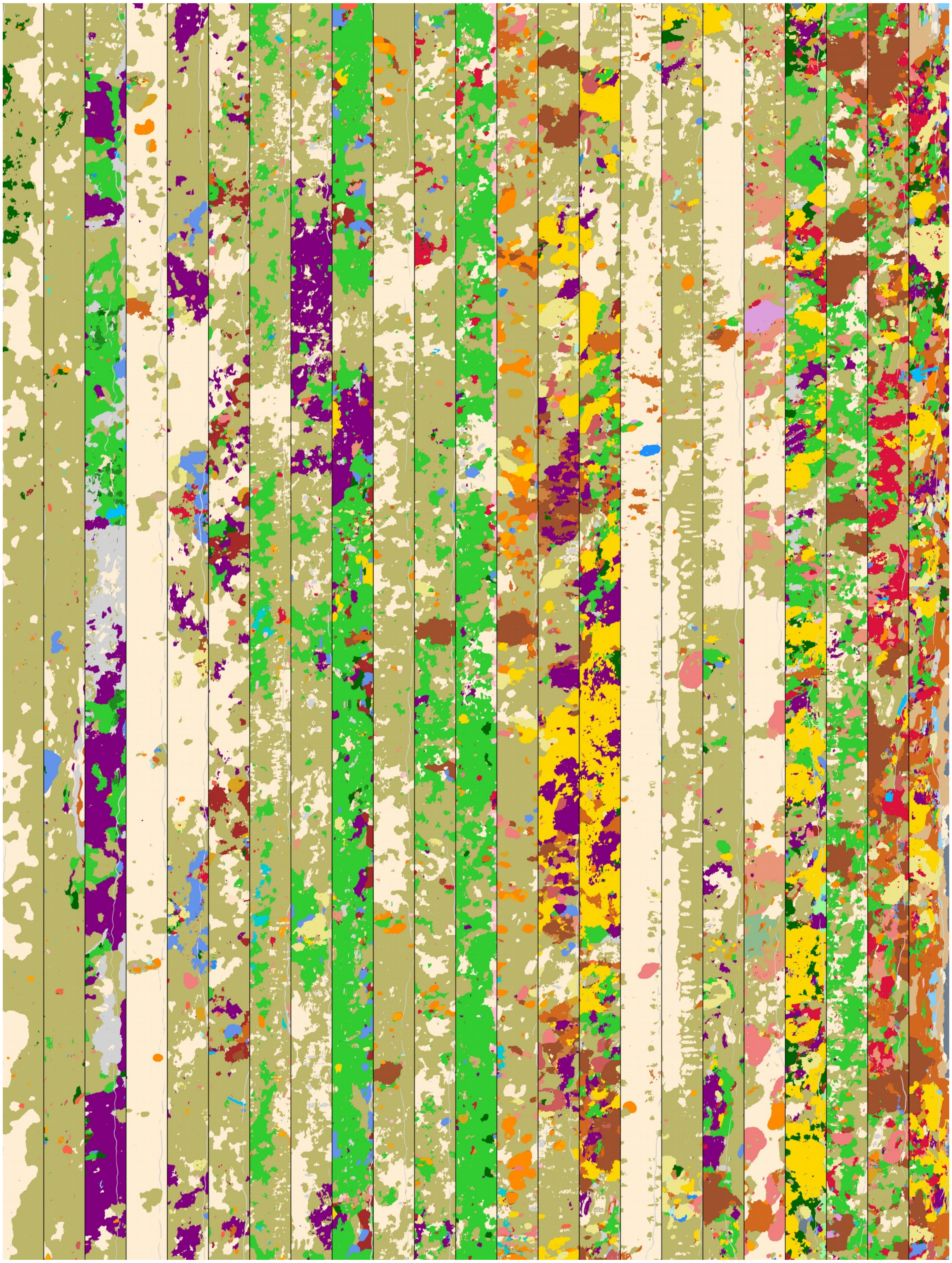

**Fig. S10.** Habitat maps montage with workflow patched+detailed+radiance. Middle section of 23 learning transects. Size of each section is 640x19920 pixels.

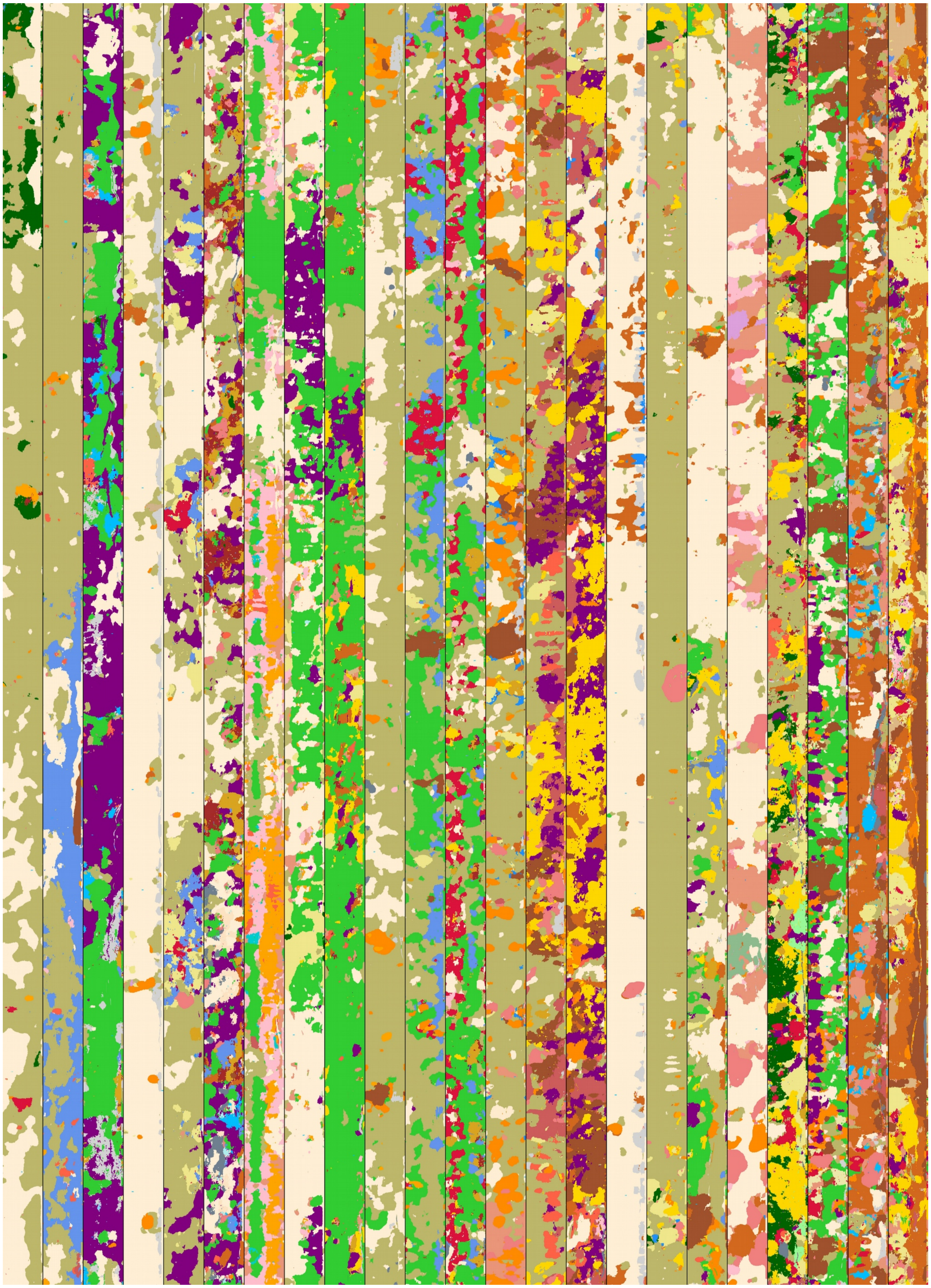

**Fig. S11.** Habitat maps with workflow segmented+detailed+reflectance. Middle section of 23 learning transects. Size of each section is 640x19920 pixels.

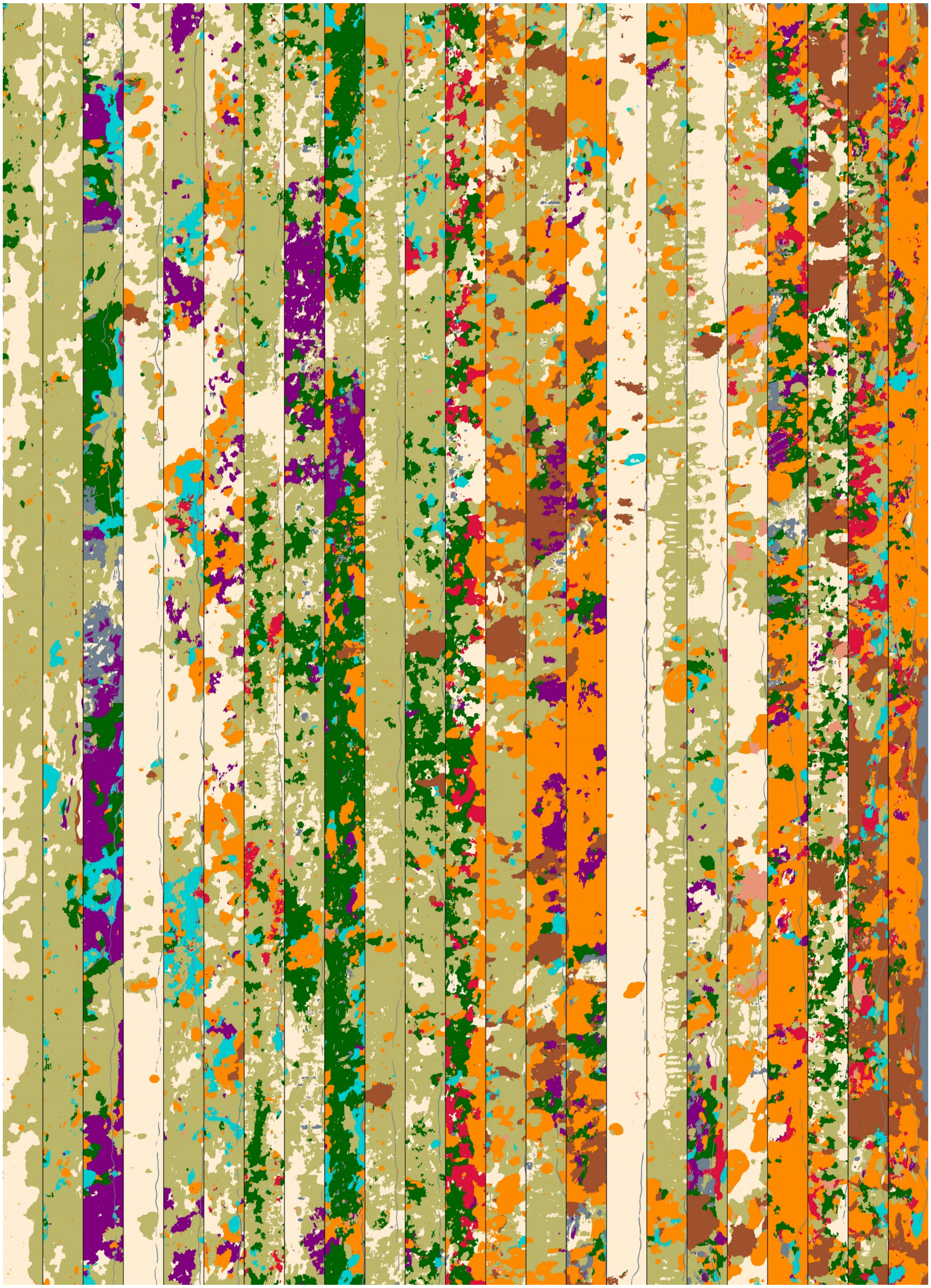

**Fig. S12.** Habitat maps montage with workflow patched + reefgroups + radiance. Middle section of 23 learning transects. Size of each section is 640x19920 pixels.

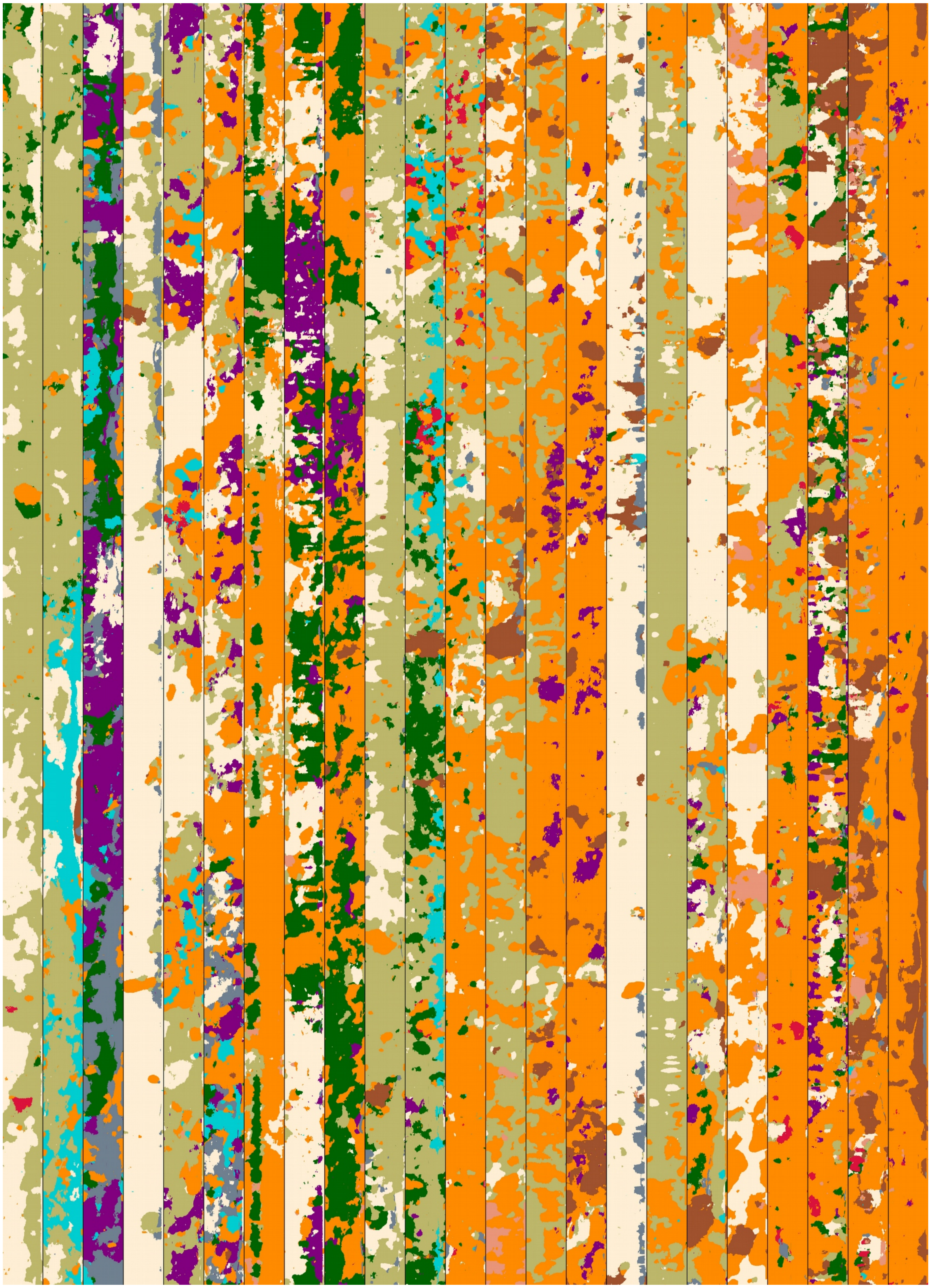

**Fig. S13.** Habitat maps montage with workflow segmented+reefgroups+reflectance. Middle section of 23 learning transects. Size of each section is 640x19920 pixels.

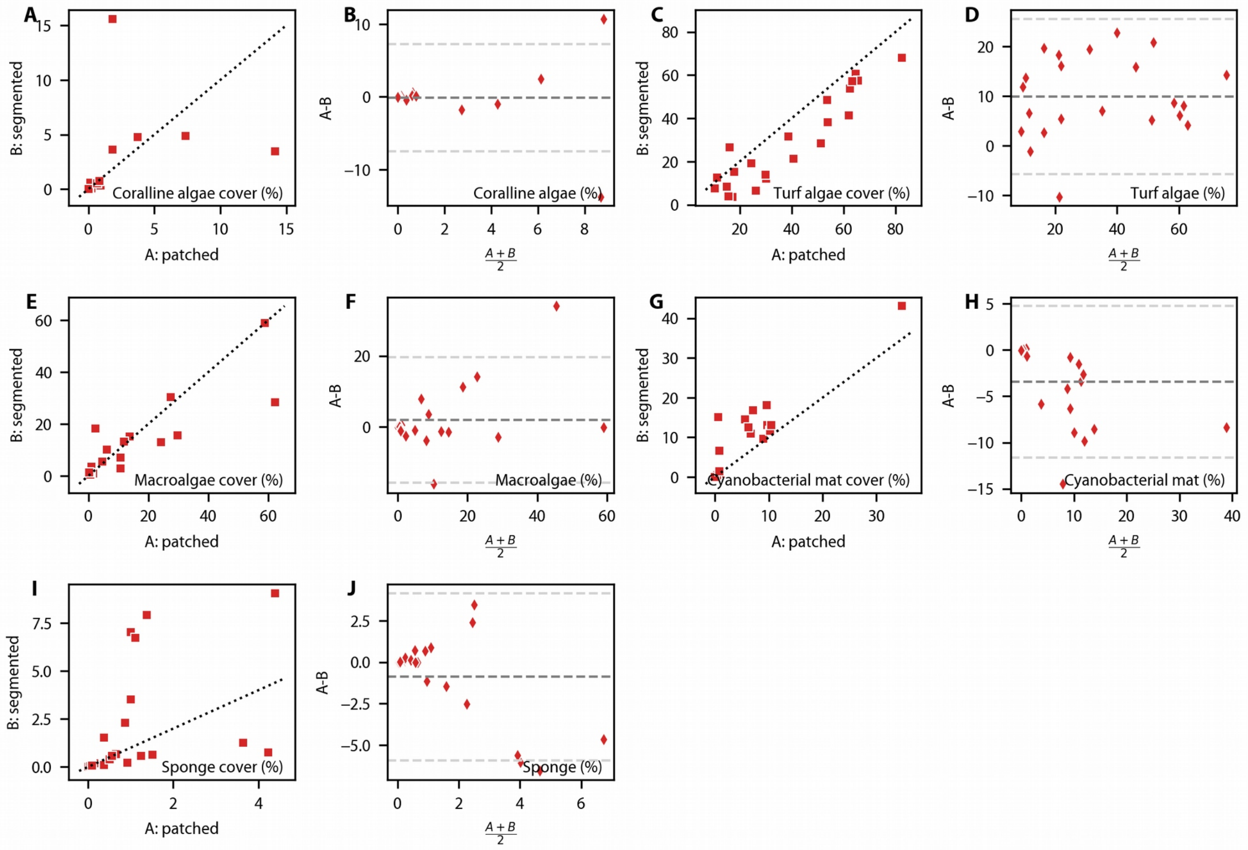


**Fig. S14.** Correlation and mean-difference plots of percent coverage from 22 transects for dominant reef classes.

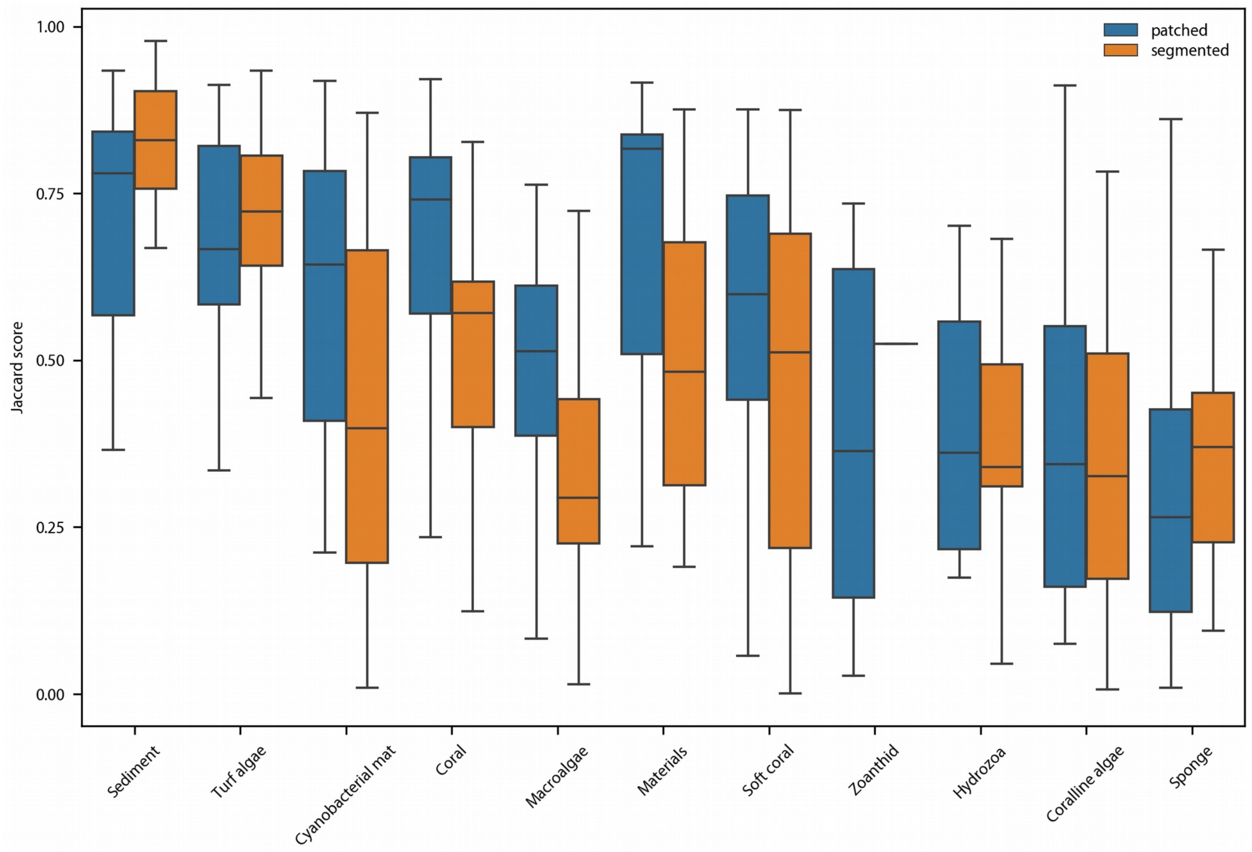


**Fig. S15.** Configuration analysis (Jaccard scores) of all 11 reefgroups labelspaces classes, expanding on panel F of Figure 6.
